# A novel rhodopsin-based voltage indicator for simultaneous two-photon optical recording with GCaMP in vivo

**DOI:** 10.1101/2024.11.15.623698

**Authors:** Vincent Villette, Shang Yang, Rosario Valenti, John J. Macklin, Jonathan Bradley, Benjamin Mathieu, Alberto Lombardini, Kaspar Podgorski, Stéphane Dieudonné, Eric R. Schreiter, Ahmed S. Abdelfattah

**Affiliations:** Institut de Biologie de l’École Normale Supérieure (IBENS), École Normale Supérieure, CNRS, INSERM, PSL Research University, Paris 75005, France; Janelia Research Campus, Howard Hughes Medical Institute, Ashburn, VA 20147, USA; Allen Institute for Neural Dynamics, Seattle, WA 98109, USA; Department of Neuroscience, Brown University, Providence, RI 02906, USA; Carney Institute for Brain Science, Brown University, Providence, RI 02906, USA

## Abstract

Genetically encoded voltage indicators (GEVIs) allow optical recording of membrane potential from targeted cells *in vivo*. However, red GEVIs that are compatible with two-photon microscopy and that can be multiplexed *in vivo* with green reporters like GCaMP, are currently lacking. To address this gap, we explored diverse rhodopsin proteins as GEVIs and engineered a novel GEVI, 2Photron, based on a rhodopsin from the green algae *Klebsormidium nitens*. 2Photron, combined with two photon ultrafast local volume excitation (ULoVE), enabled multiplexed readout of spiking and subthreshold voltage simultaneously with GCaMP calcium signals in visual cortical neurons of awake, behaving mice. These recordings revealed the cell-specific relationship of spiking and subthreshold voltage dynamics with GCaMP responses, highlighting the challenges of extracting underlying spike trains from calcium imaging.

## Introduction

Changes in membrane potential are the fundamental language of the nervous system. Although typically monitored with electrodes, less invasive visualization of voltage dynamics has come of age with genetically encoded fluorescent voltage indicators (GEVIs), allowing recordings from genetically defined neurons with exquisite spatiotemporal resolution ^1–8^. Rhodopsin domains are attractive GEVI scaffolds because of their extremely fast response times (<1 ms) to membrane potential changes. They have been successfully combined via Förster resonance energy transfer (FRET) with red-shifted fluorescent proteins or dyes for one photon (1P) *in vivo* voltage imaging^1,6^. Rhodopsin-based GEVIs, however, have historically not been used with two photon (2P) voltage imaging. Several past reports describe reduced voltage sensitivity when illuminated with 2P light9,10, but only very recent work has started to explore the causes^11,12^. In contrast, the ASAP family of GEVIs based on cpGFP and voltage-sensitive domains (VSD), have allowed 2P voltage imaging *in vivo*, thus opening access to recording from cells deeper in tissue^13,14^. However, analogous GEVIs based on red-shifted fluorescent proteins have proven much more difficult to optimize^15,16^.

Given their previous use with red-shifted fluorophores, we sought to develop an opsin-based chemigenetic GEVI that could be multiplexed with existing green fluorescent indicators *in vivo* using 2P excitation. Ideally, such a GEVI would take advantage of high-power, low-cost, fixed-wavelength lasers that output in the range 1030-1080nm, which have recently been used in fast 2P optical approaches^13,17,18^. We explored whether rhodopsin GEVIs could be suitable for this application, given their previous use with red-shifted fluorophores. We first tested a diverse set of mostly uncharacterized rhodopsin proteins for their ability to function as voltage indicators under 1P and 2P excitation. We identified novel rhodopsin domains that successfully reported membrane potential in 1P and 2P imaging. We called our best variant ‘2Photron’ and demonstrate the utility of 2Photron for 2P voltage recordings in mouse cortex and cerebellum using 1045 nm excitation. Additionally, we demonstrate simultaneous two-color 2P optical recordings *in vivo*, combining 2Photron and a red-shifted dye with either the green VSD-based GEVI JEDI-2P or the green calcium indicator GCaMP. Simultaneous calcium and voltage recording in the same neurons reveals substantial variability in the calcium indicator response to underlying action potentials that is correlated but not easily explained by parameters of the voltage signal. To our knowledge, 2Photron is the first GEVI to report membrane voltage simultaneously with GCaMP *in vivo* using 2P optical recordings.

## Results

### Exploration of diverse microbial rhodopsins as voltage indicators

We explored a diverse set of 31 microbial rhodopsin proteins, chosen to sample multiple functional rhodopsin classes and to broadly explore sequence space, as voltage indicators to identify improved GEVIs for 2P imaging (Fig. 1a and Supplementary Table 1,2). These microbial rhodopsins are cation channels, proton or sodium pumps, or have no characterized function (Fig. 1b,c). For each opsin sequence, we introduced mutations to putatively block pump or channel activity of the rhodopsin. For proton pumps, we mutated the counterion or proton donor residues from negatively charged to neutral^6,19–21^. These mutations have been shown previously to block photocurrent and generate either negative-going or positive-going GEVIs. For sodium pumps, we reasoned similar mutations would block photocurrents and generate GEVIs. For cation channels, we mutated the analogous proton donor position to a neutral residue to block channel activity^22^ and potentially generate positive-going GEVIs. We then fused each of the modified opsin sequences to HaloTag (HT) to image FRET responses of a bound fluorescent dye-HaloTag ligands (HTL)^3^, expressed them in primary rat hippocampal neurons in culture from a strong promoter and observed their expression and membrane trafficking after labeling with JF_525_-HTL (Supplementary Fig. 1 and 2).

**Fig. 1.**
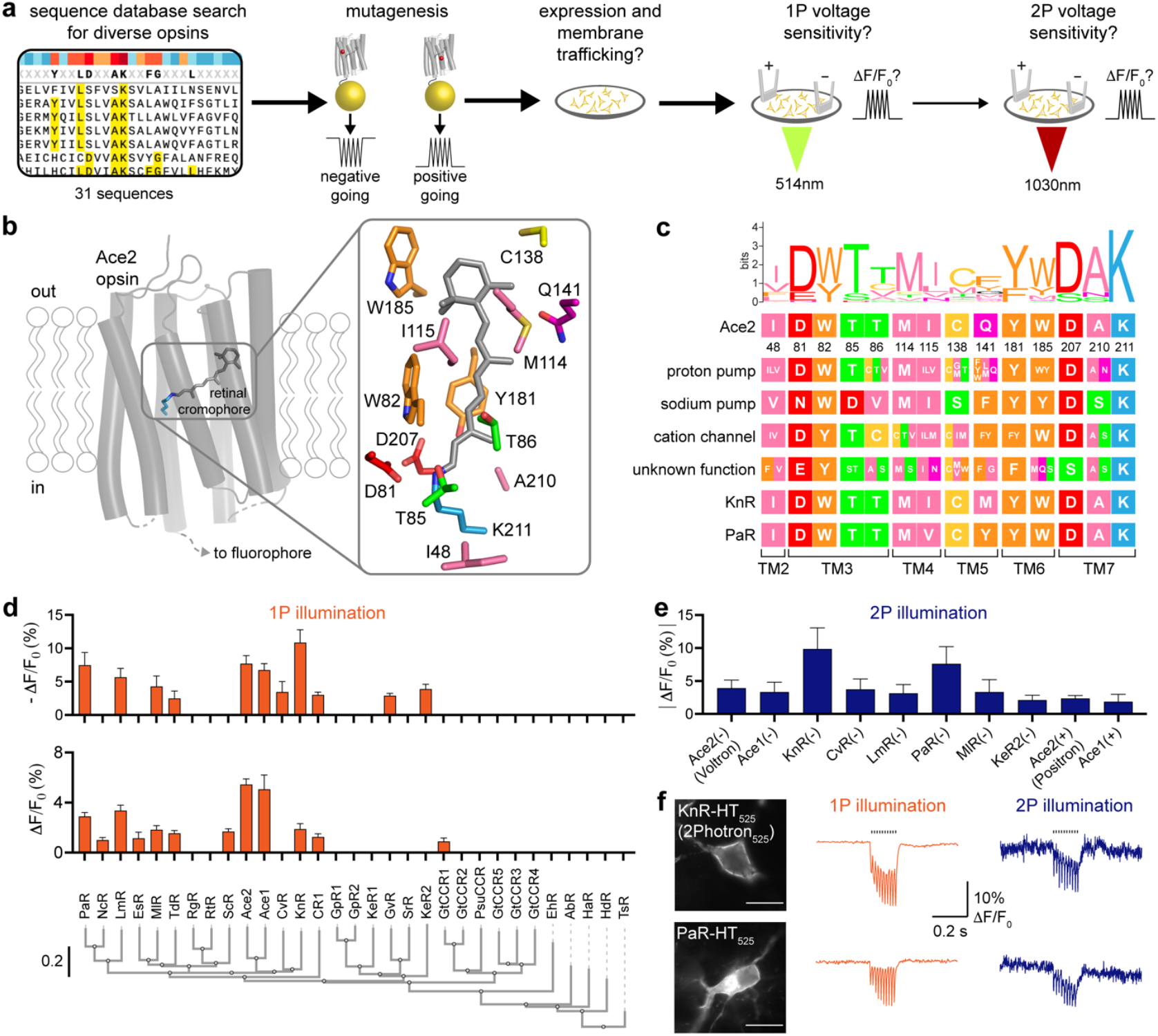
Screening diverse rhodopsin proteins for voltage sensitivity under 2P. **a**, Schematic of experimental design. Sequences of diverse opsins are aligned, homologous amino acid positions are mutated to engineer possible voltage indicator with no native pump/channel function. Opsins are then fused to HaloTag, expressed in neuron culture and labelled with JF_525_-HaloTag ligand to check for membrane trafficking, 1P voltage sensitivity and 2P voltage sensitivity. **b**, Left: cartoon of Ace2 opsin domain (PDB ID: 3AM6). Right: zoom in on retinal binding pocket in Ace2 with surrounding amino acids. Residues are colored based on their chemical properties. **c**, Comparison of key residues in the binding pocket of Ace2 opsin, different classes of rhodopsins tested for voltage sensitivity, *Klebsormidium nitens* rhodopsin (KnR), and *Podospora anserine* rhodopsin (PaR). Residues are colored according to their chemical properties as in panel b. **d**. Fluorescence response (ΔF/F_0_) of different rhodopsin-HaloTag fusions with 1P illumination (514 nm) to elicited action potentials. Cultured neurons are transfected and labeled with JF_525_-HaloTag ligand and stimulated using field potentials to elicit a train of 10 action potentials at 66Hz. Top: Fluorescence response of opsin domains with amino acid mutations to develop a negative going FRET signal. Middle: Fluorescence response of opsin domains with amino acid mutations to develop a positive going FRET signal. Bottom, phylogeny tree of the rhodopsins tested, based on opsin domain sequence alignments. The scale indicates the number of substitutions per site. **e**. Fluorescence response (ΔF/F_0_) of different rhodopsin-HaloTag fusions with 2P illumination (1030 nm) to elicited action potentials. Cultured neurons are transfected and labeled with JF_525_-HaloTag ligand and stimulated using field potentials to elicit a train of 10 action potentials at 66Hz. (-) and (+) signs indicate negative-going or positive-going version of the rhodopsin-HaloTag fusion respectively. **f**. Left: Fluorescence image of KnR-HT (2Photron) (top) and PaR-HT (bottom) labeled with JF_525_ HaloTag ligand in hippocampal neurons in culture. Scale bar: 20 µm. Middle and Right: Fluorescence response in cultured hippocampal neurons to a train of field stimuli eliciting 10 APs at 66Hz with 1P (514nm) or 2P (1030nm) illumination. Field stimulus timing is shown with black bar.

For rhodopsin variants that showed successful expression and membrane trafficking in neurons, we tested their 1P fluorescence response to membrane voltage changes by inducing a train of 10 action potentials with a field electrode (Fig. 1d and Supplementary Fig. 3). For proteins that showed strong 1P fluorescence responses to neuron stimulation, we measured their response to the same stimulation on a custom high-speed 2P microscope^18^ (Fig. 1e). We identified two rhodopsin proteins, one from the alga *Klebsormidium nitens* (KnR) and another from a fungus, *Podospora anserine* (PaR), that showed significant 1P sensitivity and retained their ability to report action potentials with 2P illumination under the conditions of our screen (Fig. 1f). PaR required grafting 39 residues from the N-terminus of the *Neurospora crassa* opsin in place of the first 44 residues, which displayed dramatically better membrane trafficking (Supplementary Fig. 4). KnR and PaR have <35% sequence identity when compared to any reported opsin-based voltage sensor (Supplementary Fig. 5).

### In vitro characterization of KnR and PaR GEVIs

To characterize the new rhodopsin GEVIs, we adapted a microscope to allow sequential imaging of the same cell under 1P and 2P illumination with simultaneous patch electrophysiology to record or stimulate. Unlike conventional 2P point scanning illumination, we created an enlarged stationary focal spot to cover a significant part of a neuron cell body. To quantify the voltage sensitivity, cultured rat neurons expressing KnR-HT_552_ or PaR-HT_552_ were held at -70 mV and then stepped to different membrane potentials using a patch electrode under sequential 1P and 2P illumination (Fig. 2a). Both KnR-HT_552_ and PaR-HT_552_ exhibited nearly identical fluorescence changes in response to voltage steps under 1P and 2P illumination (Fig. 2a,b). KnR-HT_552_ showed higher voltage sensitivity than PaR-HT_552_, giving a fluorescence change of about -24% Δ*F/F*_0_ (-24 ± 2.3 % for 1P, -24.5 ± 2.5 % for 2P) for a voltage step from -70 mV to +30 mV. Fluorescence response kinetics were also similar under 1P and 2P illumination (Fig. 2c). KnR-HT_552_ displayed fast voltage response with sub-millisecond on and off fast time constants (Supplementary Fig. 6). Both KnR-HT_552_ and PaR-HT_552_ faithfully reported action potentials as well as subthreshold voltage signals with similar fluorescence response under 1P and 2P illumination (Fig. 2d,e).

**Fig. 2.**
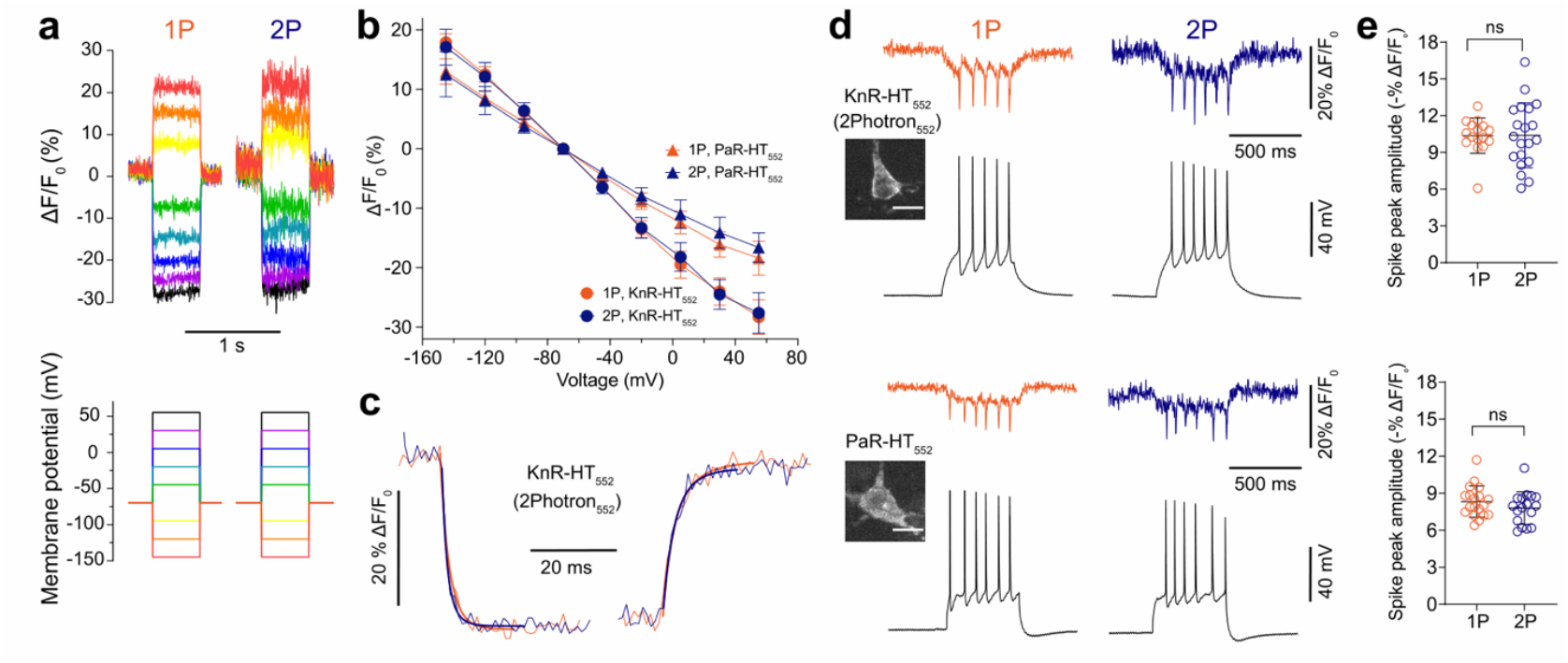
In vitro characterization of new rhodopsin GEVIs. **a**, Representative fluorescence traces of KnR-HT_552_ (top) in response to a series of 0.5 s voltage steps (bottom: from -145 mV to +55 mV in 25 mV increments) under 1P and 2P excitation. **b**, Fluorescence changes as a function of membrane voltage with KnR-HT_552_ and PaR-HT_552_ under 1P and 2P excitation. Errors bars are S.D., n = 12 (KnR-HT_552_) and 10 (PaR-HT_552_) cells. **c**, Fluorescence response of KnR-HT_552_ to a 150 mV voltage step under 1P (orange, thin line) and 2P (purple, thin line) excitation. Fluorescence traces were averaged from 5 recordings and fit using a double exponential function (thick lines). **d**, Single-trial recordings of action potentials and subthreshold membrane potentials induced by current injections in primary cultured neurons expressing KnR-HT_552_ (top) or PaR-HT_552_ (bottom) with 1P and 2P imaging at 500 Hz (colored traces) or electrophysiology (black traces). Inset: fluorescence images of primary cultured neurons expressing KnR-HT_552_ or PaR-HT_552_. Scale bar: 20 μm. **e**. Comparison of spike amplitude of KnR-HT_552_ (top) or PaR-HT_552_ (bottom) under 1P and 2P imaging at 500 Hz; n = 4 cells for KnR-HT552 and n = 3 cells for PaR-HT552; ns indicates not significant (p > 0.05). This measures only the spike amplitude, the underlying subthreshold signal is not included.

We further observed that existing rhodopsin-based voltage indicators, including Voltron, Voltron2, Positron2, and QuasAr2-HT, also exhibited similar voltage sensitivity with 1P and 2P illumination under these conditions (Supplementary Fig. 7). To explore additional 2P illumination configurations, we performed 1 kHz circular point scanning of neuron cell bodies expressing KnR-HT or Voltron2. Both KnR-HT and Voltron2 reported sub- and supra-threshold voltage changes in neurons with circular point scanning 2P excitation, with similar voltage sensitivity to 1P illumination (Supplementary Fig. 8). Collectively, we conclude that rhodopsin-based GEVIs are generally compatible with 2P imaging, at least under some illumination conditions. KnR-HT exhibited the most reliable responses under all tested 2P illumination conditions, so we named this protein 2Photron and characterized its performance further *in vivo*.

### 2P voltage recording with 2Photron in awake, behaving mice

To explore whether we could record voltage signals *in vivo* using 2P excitation, we used ultrafast local volume excitation (ULoVE) optical recording to record from individual neurons of head-fixed, awake mice free to move on a circular treadmill (Fig. 3a). We injected adeno-associated virus (AAVs) to express soma-targeted 2Photron (2Photron-ST) in mouse cortex or cerebellum. Bright, membrane-localized fluorescence of 2Photron was observed following intravenous injection of JF_552_-HTL (Fig. 3a). Using ULoVE with excitation at 1045 nm and >3.5 KHz sampling rate we recorded from 25 cells in layer II/III of the visual cortex and 19 fast-spiking cerebellar granular layer interneurons (Golgi cells) across 18 mice. We observed clear subthreshold fluctuations and negative-going spikes in single trial data with 2Photron-ST_552_, and could routinely record from individual neurons in layer II/III of visual cortex for ten minutes or more in awake mice with a mean ΔF/F for spikes of 5.72 ± 1.39% (Fig. 3b). In Golgi cells, 2Photron-ST_552_ responded to spikes with a ΔF/F of 7.88 ± 1.31% (Supplementary Fig. 9).

**Fig. 3.**
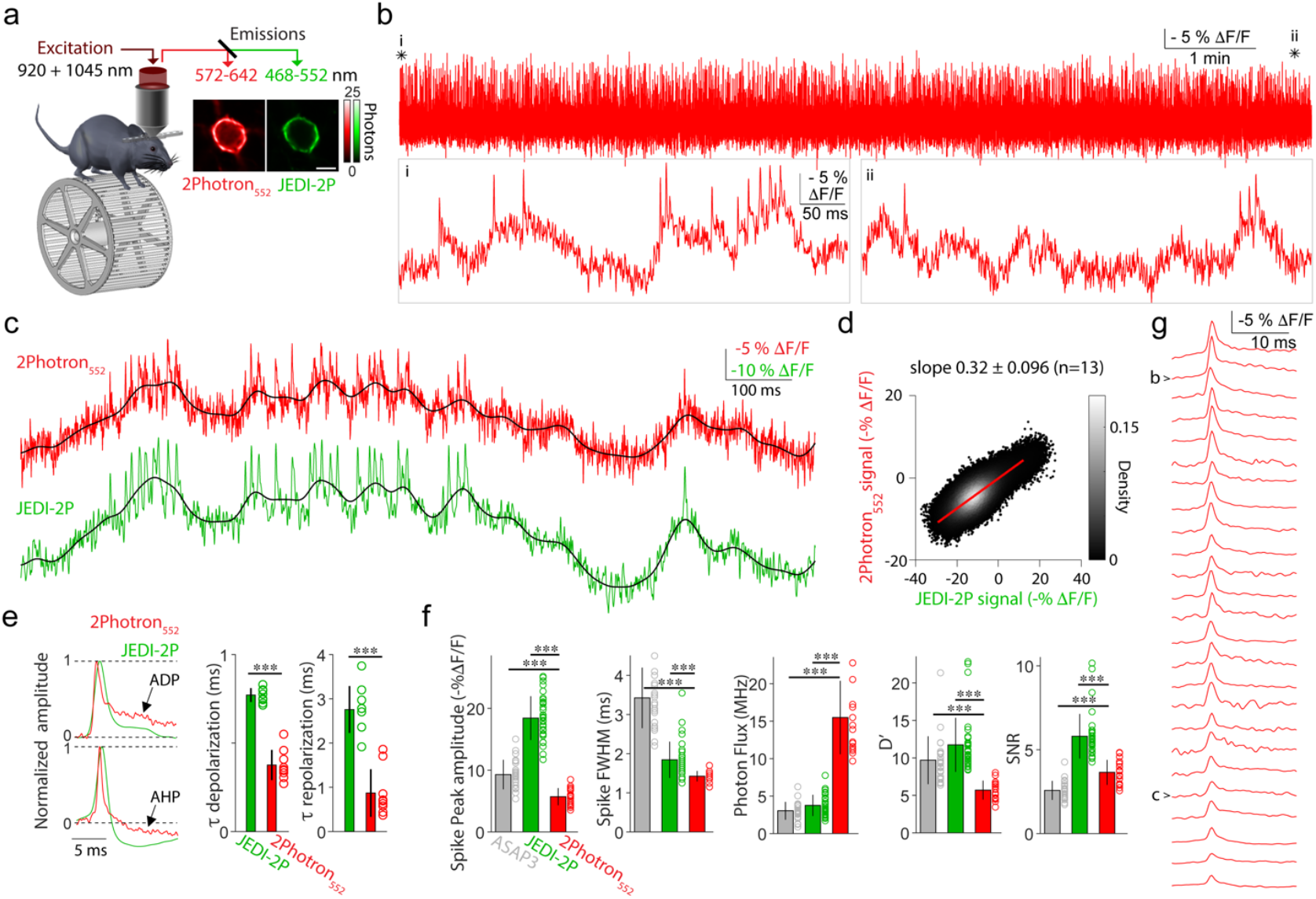
2Photron enables monitoring of sub and suprathreshold voltage in behaving mice. **a**, Left, experimental design of dual-excitation path configuration to simultaneously record 2Photron-ST_552_ and a green emission GEVI or a GECI in behaving mice. Right, fluorescence images of a cortical neuron co-expressing 2Photron-ST_552_ and JEDI-2P-Kv Scale bar: 10 μm. **b**, Representative ULoVE 3.75KHz fluorescence trace from a cortical neuron expressing 2Photron-ST_552_ in an awake, behaving mouse. Below, zoom of the trace at the beginning (i) and at the end (ii) of the recording showing the acquisition of spikes and subthreshold potentials. **c**, Representative dual recording of 2Photron-ST_552_ (top) and JEDI-2P-Kv (bottom). Black traces are low-pass filtered. **d**, Distribution of the 2Photron-ST_552_ signal as a function of the JEDI-2P-Kv signal. The data-point density is color-coded, and a linear regression is shown in red. Slope of the regression is 0.32 ± 0.096 (n=13 cells, n=4 mice). **e**, Normalized average spike waveforms from two cells co-expressing 2Photron-ST_552_ (red) and JEDI-2P-Kv (green) having an after-depolarization (top) and an after-hyperpolarization (shaded area, bottom). Quantifications are given as mean ± std for the exponential time constant of the depolarization (middle) and repolarization (right). Exponential time constant provided as mean ± std [min-max], for depolarization: 2Photron-ST_552_: 0.38 ± 0.27 ms [0.27-0.55 ms], JEDI-2P-Kv: 0.77 ± 0.04 [0.71-0.83 ms]. Wilcoxon rank sum test: p = 4.11e-5; for repolarization: 2Photron-ST_552_: 0.77 ± 0.47 ms [0.37-1.86 ms], JEDI-2P-Kv: 2.60 ± 0.39 ms [2.00-3.20 ms]. Wilcoxon rank sum test: p = 1.55e-4. The 2Photron-ST_552_ post-spike repolarization level reaches 19.60 ± 4.23 % of the JEDI post-spike repolarization level at 7.80 ± 6.92 ms. *** indicates p<0.001, data from n=9 cells, n=4 mice. **f**, Comparison of 2Photron-ST_552_ (red) with ASAP3 (grey) and JEDI-2P-Kv (green); quantifications of spike amplitude, spike FWHM, and photon flux are represented with bars ± std. *** indicates p<0.001. ASAP3 data are from Villette et al., 2019, n=23 cells. JEDI-2P-Kv data are from Liu et al., 2022, n=34 cells. 2Photron-ST_552_ (n=16 cells from 5 mice. **g**, Average single spikes from the 25 2Photron-ST_552_ recorded cells sorted top to bottom as a function of spike SNR. Example cells from Figs. 3b and c are labeled with arrowheads.

Using ULoVE with a dual-excitation path configuration^13^ (Supplementary Fig. 10), we could simultaneously record from 2Photron-ST_552_ and the GFP-based JEDI-2P-Kv^14^ in the same layer II/III visual cortical neurons following co-expression (Fig. 3a). This allowed for direct comparison of 2Photron-ST_552_ with a state-of-the-art 2P-compatible GEVI in awake mice (Fig. 3c-f), although co-expression does not allow for ideal expression of either GEVI (Supplementary Fig. 11). Distinct excitation wavelengths, acousto-optic modulator shuttering, and fluorescence emission bandpasses provided separation of the two GEVI signals (Methods and Supplementary Fig. 10). Generally, we observed good correlation between the fluorescence signals of 2Photron-ST_552_ and JEDI-2P-Kv, both for subthreshold and spike signals (Fig. 3c). A linear regression of the 2Photron-ST_552_ ΔF/F versus that of JEDI-2P-Kv for simultaneous recordings from 13 cells across 4 mice showed that JEDI-2P-Kv was ∼3x more sensitive than 2Photron-ST_552_ under these co-expression conditions (Fig. 3d). From a larger pool of 16 recorded neurons, including GCaMP co-expression, 2Photron-ST_552_ showed a mean spike amplitude of 5.72 ± 1.39 %ΔF/F, 2x lower than previously reported for JEDI-2P-Kv (from Liu et al., 2022, n=34, Wilcoxon rank sum test: p = 1.64e-8) (Fig. 3f).^14^ During spike depolarization and repolarization, 2Photron-ST_552_ exhibited 2.1x and 3.4x faster kinetics of fluorescence change than JEDI-2P-Kv (Fig. 3e). Correspondingly, spikes with 2Photron-ST_552_ were significantly narrower (Fig. 3e,f) (2Photron-ST_552_: FWHM of 1.42 ± 0.13 ms [1.15-1.70 ms], JEDI-2P: 1.84 ± 0.46 [1.25-3.55 ms] (Wilcoxon rank sum test: p = 2.91e-05, n=16 cells, 5 mice). 2Photron-ST_552_ showed brighter fluorescence output (15.5 ± 4.9 MHz [9.68-24.20 MHz], with ∼4x higher photon flux compared to JEDI-2P-Kv (3.76 ± 1.42 MHz [1.71 - 7.76 MHz], n=34, Wilcoxon rank sum test: p = 1.64e-08, Fig. 3f). Taken all together, the higher brightness of 2Photron-ST_552_ partially offsets its lower sensitivity such that the signal-to-noise ratio (SNR) for spikes is within 50% of JEDI-2P and 40% greater than ASAP3 (Fig. 3f), thus enabling single action potential detection in the red channel with a 2P microscope (Fig. 3g).

### Simultaneous GCaMP calcium and 2Photron voltage recordings in vivo

Cell-attached voltage recording together with genetically encoded calcium indicator (GECI) imaging in anesthetized rodents has shown that a given number of action potentials in a burst can generate fluorescence transients of highly variable amplitudes^23–26^. This variability cannot be explained by a unified biophysical model applying to all cells^27^ and could partly be due to the perturbation of the recording electrode. Using AAVs, we co-expressed 2Photron-ST_552_ with GCaMP6/8f (Fig. 4a-e and Supplementary Fig. 11f) to probe with minimal invasiveness the relationship between neuron electrical activity and cytoplasmic calcium transients in layer II/III of the neocortex of awake mice.

**Fig. 4.**
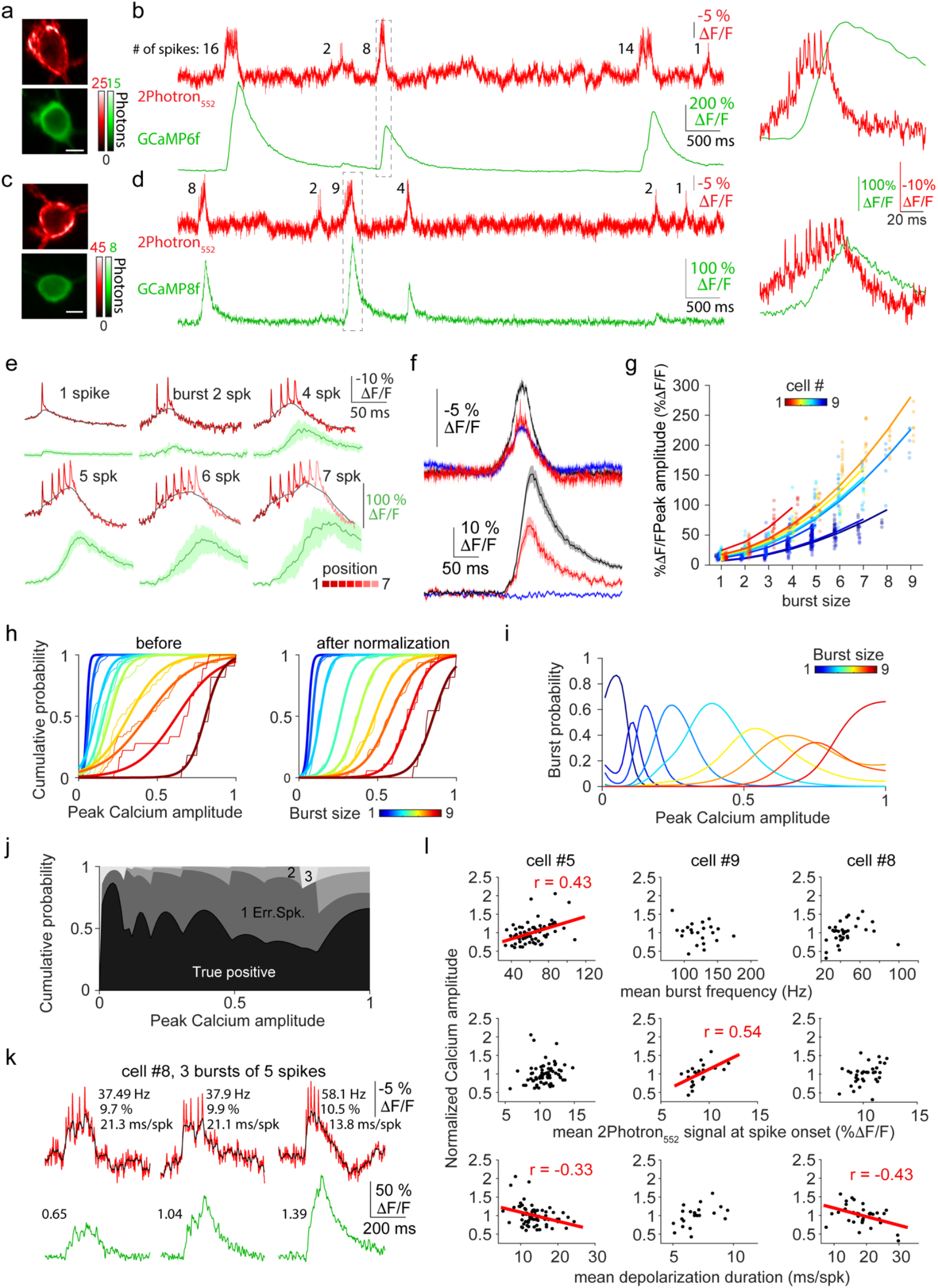
Multiplexed 2Photron-ST_552_ and GCaMP recordings reveal contributions from spiking and subthreshold voltage dynamics to calcium transient amplitudes. **a**, Single 2P-image planes showing 2Photron-ST_552_ (top) and GCaMP6f (bottom) expression in mouse cortex, scale bar 5 µm. **b**, Representative multiplexed 2Photron-ST_552_ (top) and GCaMP6f (bottom) recording. Right, zoom of recording within the dashed box encompassing a burst. **c**,**d**, same as **(a)** and **(b)** but using GCaMP8f. **e**, Average bursts where each 2Photron-ST_552_ spike (red trace) is a mean of spikes within the burst. Black trace, drifting average MLspike baseline spike trigger signal. Green trace, average calcium transient. Filled area is mean ± std. **f**, Top, 2Photron-ST_552_ signal for depolarizations without spikes (blue, n=119), depolarizations with spikes (black, n=164), and depolarizations with spikes selected to match subthreshold depolarizations (red n=52). Bottom, corresponding GCaMP8f signals, without spikes: 0.03 ± 0.46 % ΔF/F, with spikes: 23.2 ± 0.18 % ΔF/F. **g**, Distributions of GCaMP fluorescence amplitudes as a function of spike burst size for individual cells, color-coded by the scaling factor obtained from quadratic fits. **h**, Cumulative distributions and sigmoid fit of GCaMP8f calcium transient amplitudes as a function of burst sizes. Left, before normalization. Right, after normalization. **i**, Probability distribution of burst occurrence as a function of the normalized amplitude. **j**, Cumulative spike distribution as a function of the normalized amplitude. Black, true positive. Grey: false positive, ranging in shades of grey from ± 1 to 4 erroneous spikes. **k**, Three example bursts of five spikes from the same neuron (2Photron-ST_552_, top), highlighting variation in GCaMP fluorescence response (bottom). Indicated values for voltage traces are mean intraburst firing frequency, average 2Photron-ST_552_ signal at burst onset, and burst duration. GCaMP responses sorted according to peak amplitude, noted as a fraction of the mean response for this cell and spike burst size. **l**, Normalized calcium transient peak amplitudes as a function of the three voltage-dependent metrics measured in **(k)**. Lines represent significant correlations with the relative amplitude of the GCaMP signal.

We extracted 1686 bursts of spikes from simultaneous voltage and calcium recordings obtained from 20 cells in 6 mice. We first verified that sub-threshold depolarizations without spikes did not evoke detectable GECI (GCaMP8f) fluorescence changes, while amplitude-matched depolarizations with spikes did (Fig. 4f). We then observed, as previously reported, a supralinear GCaMP ΔF/F increase as a function of burst size that was well fitted by a quadratic function (Fig. 4g and Supplementary Fig. 12a). The range of GCaMP ΔF/F response amplitudes, however, displayed large cell-to-cell variability that could be quantified by the amplitude scaling factor of the quadratic fit of each cell (mean ± std: 14.2 ± 5.34, range: 6.9-24.4, n=9 cells). Normalizing GCaMP fluorescence amplitudes by this fitted scaling factor for each cell nearly abolished the intercellular source of variability; it reduced the average CV of the population amplitude distribution for a given spike number from 0.41 ± 0.13 to 0.26 ± 0.08 (Supplementary Fig. 12b), which is similar to the average CV calculated from each individual cell (0.21 ± 0.06, rank sum test, p=0.11).

The probability distribution of spike burst size for a given GCaMP response amplitude could then be calculated for the scaled data, yielding a best-case situation to test for burst size prediction (Fig. 4i). Because of the overlap between these probability distributions, prediction of spike burst size for a given GCaMP response amplitude has significant uncertainty. The percentage of GECI transients for which the most likely spike number matched the GEVI recording was 51.7% from raw data and improved to 60.5% after normalization (Fig. 4j). To provide a quantitative estimate of the spike assignment error, we computed the absolute value of the difference between the most likely spike number and the actual spike number. We found that normalization decreased the error from 0.517 spikes to 0.415 spikes per event. These results indicate that the best estimation, knowing the true distribution of GCaMP response amplitudes, would still lead to an approximate determination of spike number per transient. Using a realistic normalization for GECI-only recordings (Supplementary Fig. 12c-e), where each cell’s transient amplitude is normalized by the highest amplitude seen for that cell, we calculated an intermediate true spike number estimation of 55.1% and 0.591 average spike-number error per event. Replication of this analysis using a dataset from the literature^28^, which combined *in vivo* juxtacellular electrophysiology and GCaMP8f calcium imaging, yielded similar results (Supplementary Fig. 12f-h), indicating that GCaMP response variability is not a consequence of voltage recording conditions.

GCaMP response amplitude variability for a given number of spikes might be explained both by the biophysics of calcium binding to the GECI^27^ and by the time and voltage dependence of the various sources of calcium influx. We examined whether the variability of GCaMP response around its mean (for the same cell and the same spike burst size) was correlated to any parameters of the voltage signal during the burst. We found that the 2Photron-ST_552_ signal preceding the last spike of the burst increased with burst size from 6.61 to 10.62 % ΔF/F and then saturated (99th percentile after 5.01 spikes; Supplementary Fig. 12i). We thus measured 1) the mean 2Photron-ST_552_ signal at spike onset during the burst, 2) the mean intra-burst frequency, and 3) the mean duration of the depolarization plateau underlying each burst to see whether they could explain the GCaMP response variability. We found that, in 7 out of 9 cells, GCaMP response amplitude variation correlated significantly with one of these three parameters (Pearson coeffcient range: -0.434 to 0.54, Figure 4k, l) while none of the cells displayed significant correlation of the GCaMP response amplitude with the interval from the preceding burst (Pearson coeffcient range: -0.222 to 0.2718, Supplementary Fig. 12j). As these three parameters were either anti-correlated (10/27 pairs, Pearson coeffcient range: -0.861 to -0.549) or correlated (9/27 pairs, Pearson coeffcient range: 0.311 to 0.465) for the nine neurons recorded, we additionally performed partial correlation analysis. Notably, in 6/9 cells, GCaMP response amplitudes displayed a significant partial correlation with either the mean intra-burst frequency (2/6 cells R: -0.239 & 0.3824), the mean GEVI signal at spike onset (5/6 cells, R: -0.35 to 0.737) or the depolarization plateau underlying each burst (4/6 cells, R: -0.423 to 0.666). These data taken together suggest that the relationship between the GCaMP response amplitudes and electrophysiological parameters of a spike burst is multifactorial and cell-specific, making precise inference of spikes from the GCaMP ΔF/F signal difficult.

## Discussion

Despite an increasing number of reports describing application of novel and improved GEVIs for biology, optically recording voltage remains challenging, largely due to the demands imposed by rapid acquisition of fast signals like action potentials. Recent high-speed 2P microscopy approaches have taken advantage of the high power output, good stability, and low cost of fixed wavelength lasers that output in the range of 1030-1080nm^17,18^. These wavelengths, however, do not efficiently excite GFP-based reporters^29^. Here we used a chemigenetic approach to combine a novel rhodopsin voltage sensor domain with fluorescence output from bright synthetic dyes, such as JF_552_, that can be efficiently excited beyond 1030nm and have a red-shifted emission that is separable from GFP for multiplexed recordings.

Although past efforts to image rhodopsin-based GEVIs with 2P largely failed, two successful efforts in addition to our work here have recently appeared as preprints^11,12^. Cumulatively, these efforts shed significant light on the problem. A common conclusion is that FRET rhodopsin GEVIs can indeed be used with 2P illumination, but the details of the illumination conditions can influence the amplitude and kinetics of fluorescence change^11^. It remains unclear from which intermediates (or ground state) of the photocycle the voltage sensitivity of rhodopsin arises. Different illumination schemes (e.g. wavelength, frequency and intensity etc.) might drive the rhodopsin into different photointermediate population distributions, potentially affecting its voltage sensitivity. In our hands, parallel, continuous 2P illumination performed well, and circular point scanning also worked for *in vitro* applications (Supplementary Table 3). ULoVE excitation from 80 MHz laser pulses allowed good quality *in vivo* optical voltage recordings from 2Photron (Supplementary Table 3).

In this work, we engineer a new rhodopsin GEVI, 2Photron, and demonstrate *in vivo* 2P voltage recording with single-trial, single-spike resolution over many minutes, compatible with a variety of behavioral paradigms in mice. Using a red-shifted dye with 2Photron frees the commonly used green fluorescence channel for simultaneous multiplexed monitoring of GFP-based indicators, and allowed us to perform simultaneous recordings of voltage and calcium using 2Photron and GCaMP. Our results highlighted the difficulty of precisely assigning spike numbers and timing using only calcium imaging, due to cell-to-cell variability in the voltage to calcium relationship that could not be accounted for using only GCaMP data. Voltage imaging will likely play an increasingly important role in reading out action potentials from circuits *in vivo*. Multiplexed 2P recordings will enable further dissection of circuit function through correlation of voltage signals with neurotransmitters, neuromodulators, or other aspects of cell signaling.

There are several reasons to be optimistic about the future of voltage imaging *in vivo*. This and other recent work show that it is possible to use rhodopsin-based GEVIs for 2P recordings in animals, substantially expanding the sensor options available. Future refinement of 2P illumination parameters, perhaps resulting from deeper characterization of photophysical properties, could enable higher SNR imaging with rhodopsin GEVIs. Additionally, we performed no optimization of the voltage sensitivity of 2Photron beyond grafting a mutation from the literature to block proton pumping by the *Klebsormidium* rhodopsin. Thus, there could be significant headroom for improvement of the sensitivity of 2Photron via mutagenesis. Screening for performance using 2P illumination that mimic *in vivo* conditions should generate better rhodopsin GEVI performance.

## Acknowledgements

We acknowledge the Howard Hughes Medical Institute and the Janelia Visiting Scientist Program for support. We thank Janelia shared support teams, especially Deepika Walpita and Kim Ritola, for experimental support. We thank Luke Lavis for help and advice on fluorescent dyes. We thank Boaz Moffar and Lin Zhong for help and advice on *in vitro* 2p imaging. We thank the IBENS imaging facility (IMACHEM-IBiSA), member of the French National Research Infrastructure France-BioImaging (ANR-10-INBS-04), which received support from the “Fédération pour la Recherche sur le Cerveau—Rotary International France” (2011) and from the program «Investissements d’Avenir» ANR-10-LABX-54 MEMOLIFE. We thank Caroline Mailhes Hamon for helping with surgeries and the IBENS PFL2 platform, IBENS animal facility, and Y. Cabirou for custom mechanical production. We thank Annick Ayon and Walther Akemann for helpful discussions. The project was supported by the Searle Scholar Program (A.S.A), NIH New innovator award 1DP2MH129956 (A.S.A), NIH BRAIN Initiative grant 1U01NS103464 (S.D.), the ANR (MuVRICC, AAPG JCJC, ANR-20-CE16-0018-01), INSERM (J.B., B.M. and S.D.); CNRS (V.V.); and ENS (J.B., V.V., A.L. and S.D.).

