## Supplementary Figures and Tables for "A novel rhodopsin-based voltage indicator for simultaneous two-photon optical recording with GCaMP in vivo"

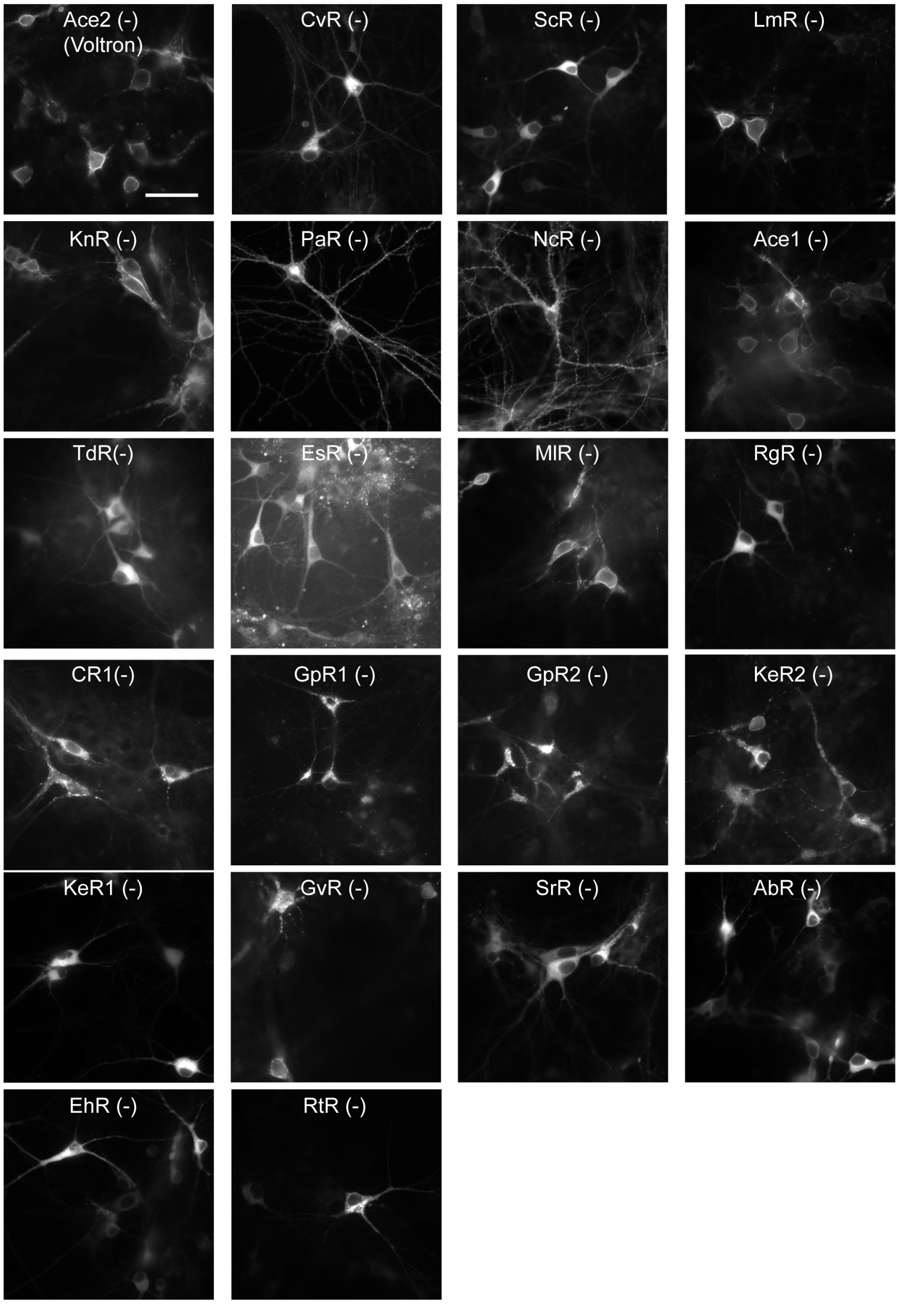


**Supplementary Figure 1. Expression of negative-going voltage sensors under 1P illumination.** Fluorescence images of hippocampal neurons in culture expressing rhodopsin-HaloTag fusions with point mutations to generate negative-going voltage sensors. Scale bar: 50 μm.

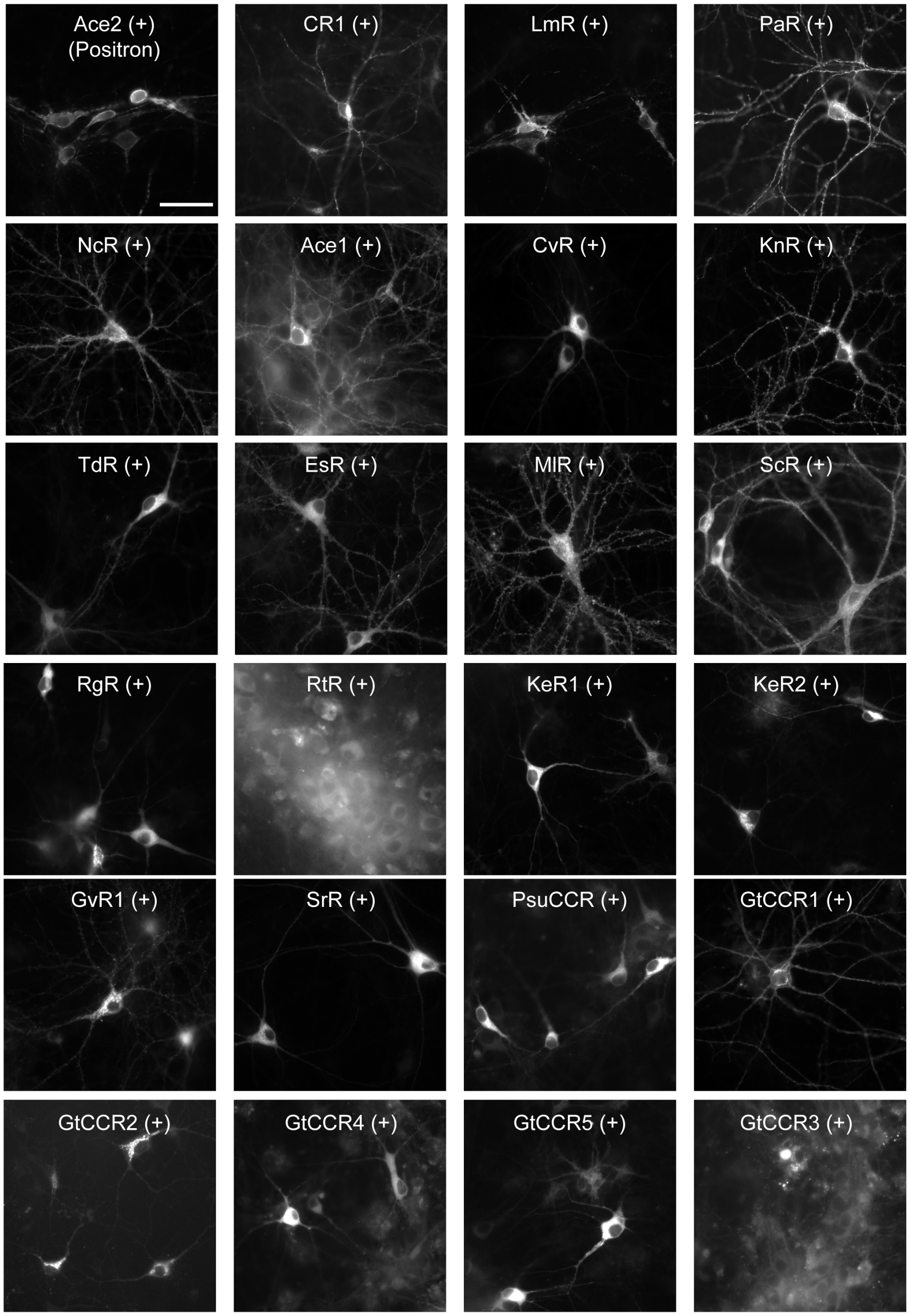


**Supplementary Figure 2. Expression of positive-going voltage sensors under 1P illumination.** Fluorescence images of hippocampal neurons in culture expressing rhodopsin-HaloTag fusions with point mutations to generate negative-going voltage sensors. Scale bar: 50 μm.

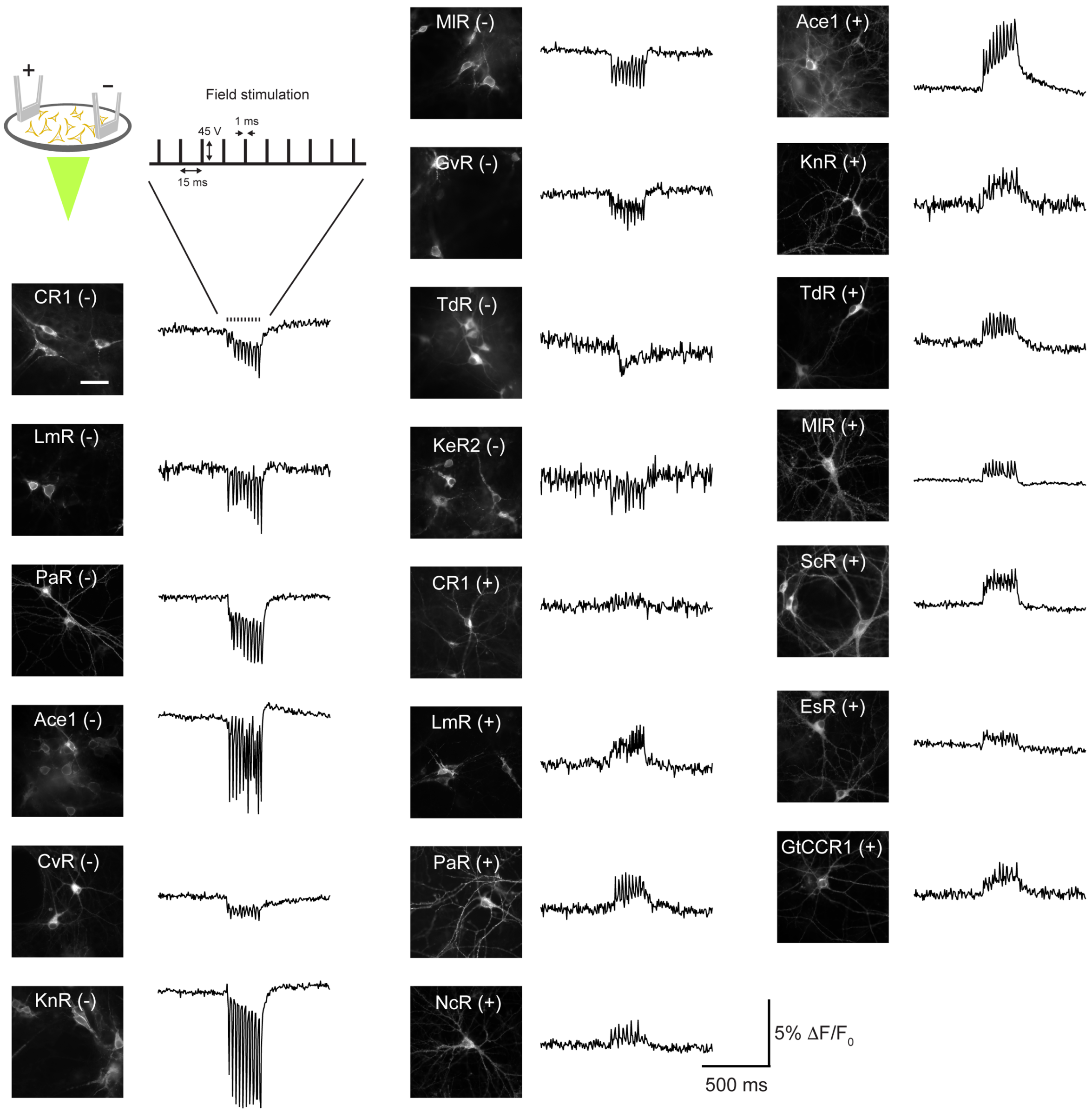


**Supplementary Figure 3. Example fluorescence response of rhodopsin-HaloTag fusions that exhibited fluorescence change to field stimulation. Top Left:** Schematic of field stimulation protocol. **Left:** Fluorescence images of hippocampal neurons in culture expressing rhodopsin-HaloTag fusions. Scale bar: 50 μm. Right: Fluorescence response to 66Hz field stimulation.

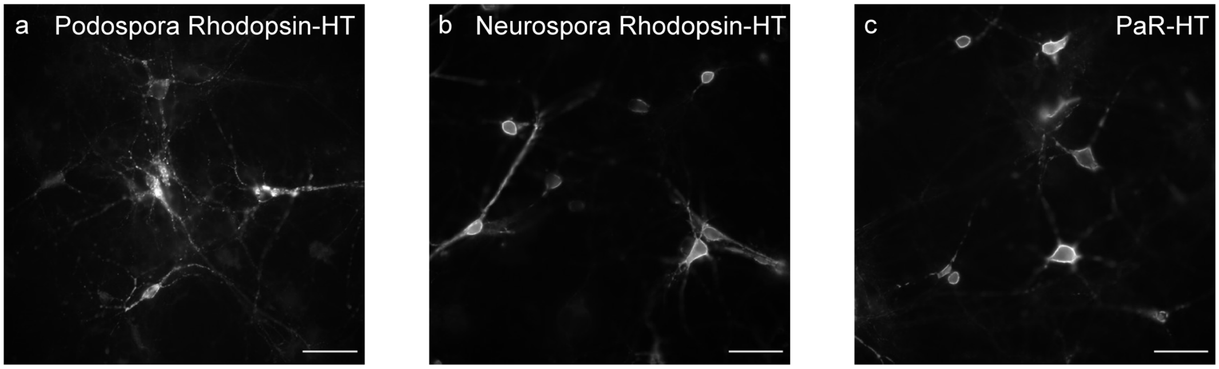


**Supplementary Figure 4. Trafficking of Podospora and Neurospora rhodopsin domains fused to HaloTag in neuron culture. a**, *Podospora anserine* rhodopsin when fused with HaloTag and labeled with JF525 displayed poor membrane trafficking in neuron culture. **b**, *Neurospora crassa* rhodopsin when fused with HaloTag and labeled with JF525 displayed good membrane trafficking in neuron culture. **c**, Replacing the N-terminal 44 residues of Podospora rhodopsin with the N-terminal 39 residues from Neurospora rhodopsin improved better membrane trafficking. Scale bar: 40 μm.

>KnR-HT (2Photron)
MVYSHADAAGWTLYWITYGIMAVTALIFFAMSLRRPIQQRSHHYTSFLIVAIASLAYYAMASQG
GNTRIRVYQGPADSYRQIFWARYVNWFFTTPLLLLDLVLLSNLSKLRIAAIMVADIFMILTGLF
GAVEARSNKWGWFVFGCIFMLYIFYELLVNVRKGAYARGGQHGMLYSVLLVWLLILWVQYPVVW
GLAEGSSTVSSDTEIAWYAALDICAKCVFGFILLLGIESIDRKRIGTGFPFDPHYVEVLGERMH
YVDVGPRDGTPVLFLHGNPTSSYVWRNIIPHVAPTHRCIAPDLIGMGKSDKPDLGYFFDDHVRF
MDAFIEALGLEEVVLVIHDWGSALGFHWAKRNPERVKGIAFMEFIRPIPTWDEWPEFARETFQA
FRTTDVGRKLIIDQNVFIEGTLPMGVVRPLTEVEMDHYREPFLNPVDREPLWRFPNELPIAGEP
ANIVALVEEYMDWLHQSPVPKLLFWGTPGVLIPPAEAARLAKSLPNCKAVDIGPGLNLLQEDNP
DLIGSEIARWLSTLEISGEPTTKSRITSEGEYIPLDQIDINVFCYENEVQSQPILNTKEMAPQS
KPPEELEMSSMPSPVAPLPARTEGVIDMRSMSSIDSFISCATDFPEATRF

Klebsormidium nitens opsin
HaloTag
membrane targeting
ER export
soma targeting

>PaR-HT
MIHPEQVADMLRPTTSTTSSHVPGPVPTVVPTPTEYQTLGETGHRTLWVVFALMVLSSGFFAFM
SWNVPISKRLYHVITTLITITASLSYFAMASGHVTSFSCTPAKDHHKHVPDVGYTECRQVFWGR
YVNWAITTPLLLLDLSLLAGIDGAHTLMAVIADVIMVLSGLFASQGETATQRWGWYAIGCVSYL
FVIWHVALHGARTVTAKGRGVTRLFSSLALFTFVLWTAYPIVWGIADGAHRTTVDTEILIYAVL
DILAKPVFGLWLLFSHRSLAETNIGTGFPFDPHYVEVLGERMHYVDVGPRDGTPVLFLHGNPTS
SYVWRNIIPHVAPTHRCIAPDLIGMGKSDKPDLGYFFDDHVRFMDAFIEALGLEEVVLVIHDWG
SALGFHWAKRNPERVKGIAFMEFIRPIPTWDEWPEFARETFQAFRTTDVGRKLIIDQNVFIEGT
LPMGVVRPLTEVEMDHYREPFLNPVDREPLWRFPNELPIAGEPANIVALVEEYMDWLHQSPVPK
LLFWGTPGVLIPPAEAARLAKSLPNCKAVDIGPGLNLLQEDNPDLIGSEIARWLSTLEISGEPT
TKSRITSEGEYIPLDQIDINVFCYENEVQSQPILNTKEMAPQSKPPEELEMSSMPSPVAPLPAR
TEGVIDMRSMSSIDSFISCATDFPEATRF

Podospora anserine opsin (underlined portion from N-terminus of Neurospora crassa opsin)
HaloTag
membrane targeting
ER export
soma targeting

**Supplementary Figure 5. Amino acid sequence of KnR-HT (Top) and PaR-HT (bottom), with sequence features annotated.**

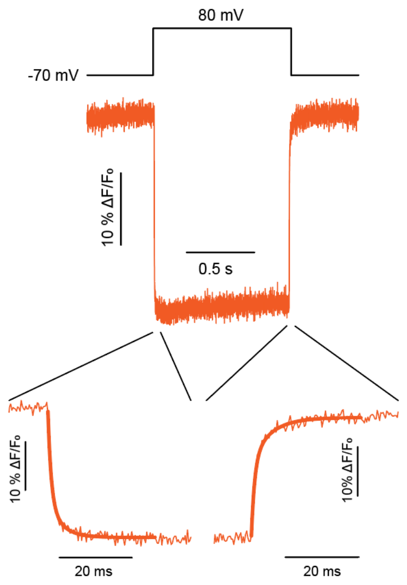


| | Activation (-70 to 80 mV) | Deactivation (80 mV to -70 mV) |
|---|---|---|
| $\tau_{fast}$ (ms) | 0.75 ± 0.15 | 0.91 ± 0.15 |
| $\tau_{slow}$ (ms) | 3.45 ± 0.74 | 4.11 ± 0.89 |
| % fast | 67 ± 1.9 | 59.3 ± 4.9 |

**Supplementary Figure 6. Characterization of the kinetic properties of KnR-HT (2Photron) under 1P illumination.** Left: Representative fluorescence response of 2Photron labeled with JF552-HTL in a cultured neuron to a 150 mV potential step delivered in voltage clamp. Insets: Zoom in on change of 2Photron fluorescence to depolarization and hyperpolarization. Solid line is double exponential fit according to $\Delta F/F(t) = A1{*}e^{-t/\tau fast} + A2{*}e^{-t/\tau slow}$. Image acquisition rate was 3.2 kHz. Right: Quantification of 2Photron kinetics in primary neurons in culture. Errors are S.D. n = 6 cells.

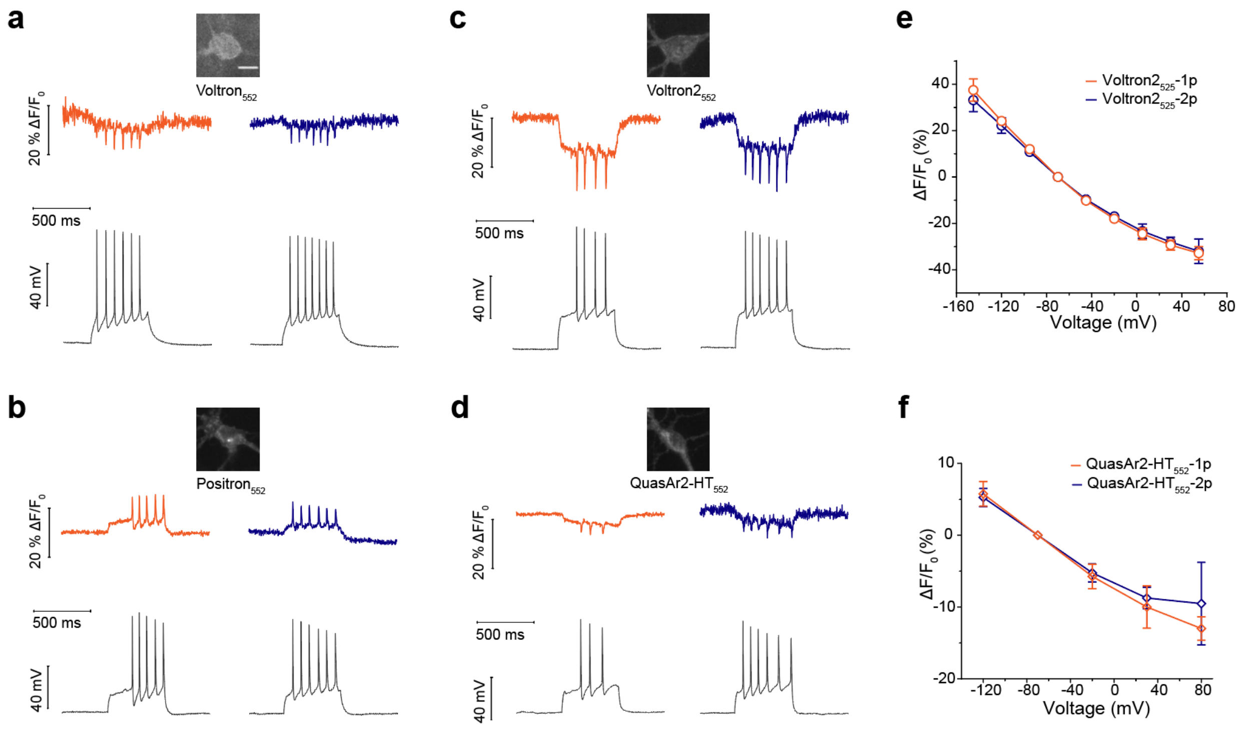


**Supplementary Figure 7. Rhodopsin derived GEVIs are generally compatible with 2p excitation.** Single-trial recordings of action potentials and subthreshold voltage signals from current injections in cultured primary rat hippocampal neurons expressing Voltron (**a**), Positron2 (**b**), Voltron2 (**c**) or QuasAr2-HT (**d**) and labeled with JF552-HTL, using 500-Hz imaging (top, fluorescence) or electrophysiology (bottom, membrane potential) under sequential 1p and stationary 2p illumination. Fluorescence changes as a function of membrane voltage with Voltron2 (**e**) and QuasAr2 (**f**) under 1p and 2p light excitation. Scale bar: 20 µm. Errors are S.D. n = 7 (**e**) or 5 (**f**) cells.

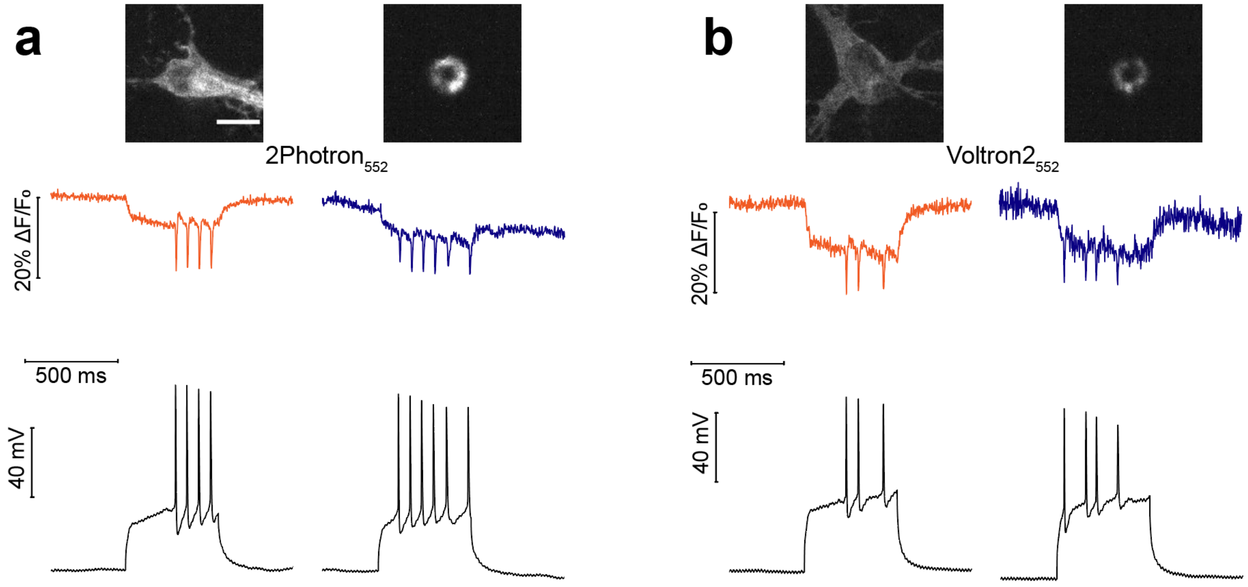


**Supplementary Figure 8. KnR-HT and Voltron2 are compatible with point scanning 2p excitation.** Single-trial recordings of action potentials and subthreshold voltage signals from current injections in cultured primary rat hippocampal neurons expressing 2Photron (**a**) and Voltron2 (**b**) and labeled with JF552-HTL, using 500 Hz imaging (top and middle: fluorescence) or electrophysiology (bottom: membrane potential) under sequential 1P (left) and circular point scanning 2P illumination (right). Scale bar: 20 μm.

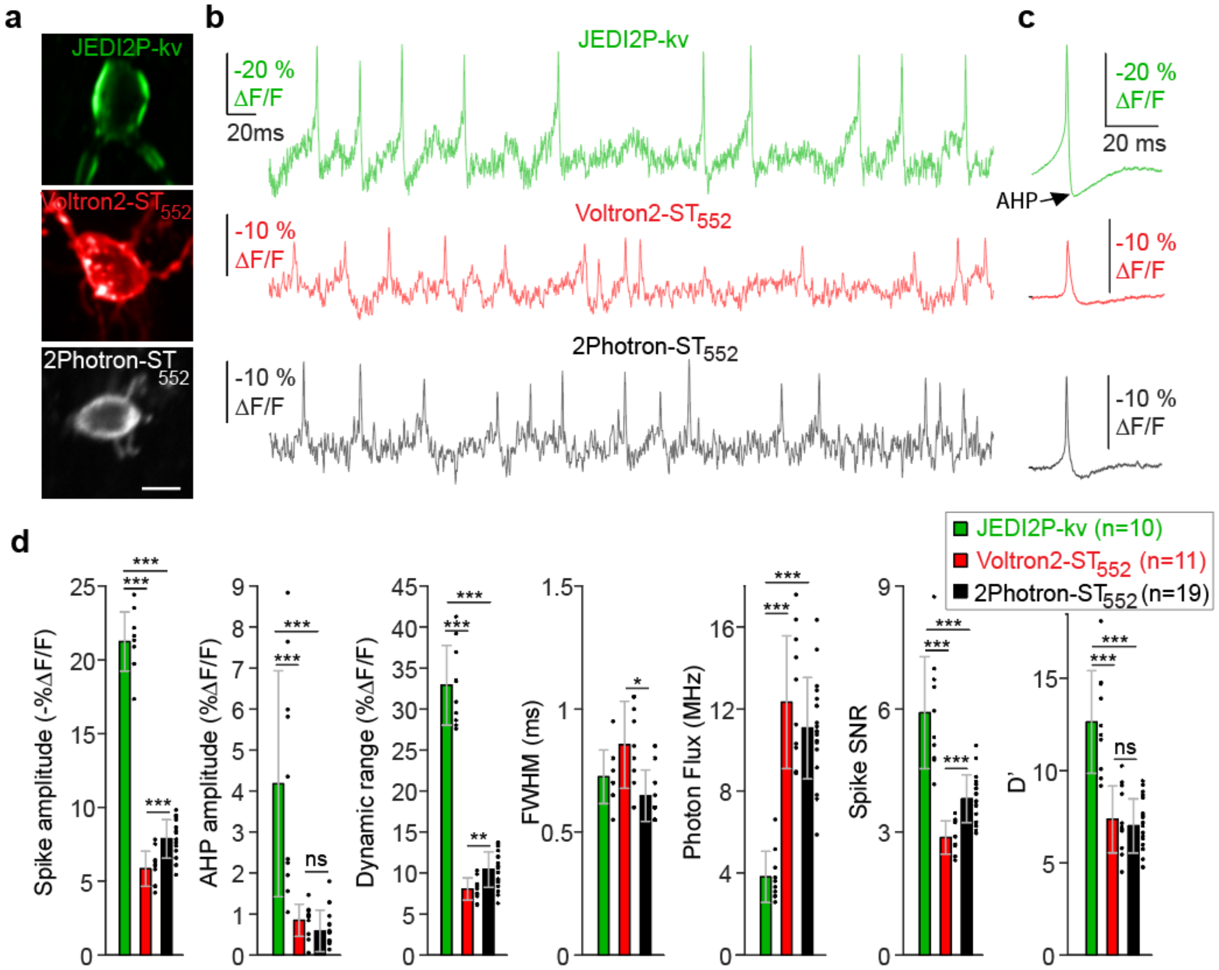


**Supplementary Figure 9. 2Photron-ST$_{552}$ outperforms Voltron2-ST$_{552}$ recording from fast-spiking cerebellar granular layer glycinergic interneurons *in vivo*. a**, Representative single 2P-image planes of GEVI expression in cerebellar Glyt2 positive granular layer neurons *in vivo*. Top, JEDI2P. Middle, Voltron2-ST$_{552}$. Bottom, 2Photron-ST$_{552}$. **b**, ULoVE recordings from cells in A at >3.5KHz in awake-behaving mice. **c**, Average spike waveforms from cells in B (mean ± sem). Note the depolarization ramp preceding and the afterhyperpolarization (AHP) following the spike. **d**, Comparisons with the state-of-the-art GEVIs. Quantifications of spike amplitude, afterhyperpolarization potential (AHP), the dynamic range, spike FWHM, the recorded photon flux, the spike Signal to Noise ratio (SNR), and the spike detectability index (D’). Green, JEDI-2P-Kv, n=10, Red, Voltron2-$ST_{552}$, n=11. Black, 2Photron-$ST_{552}$ n=19. ± std, *** indicates $p<0.001$, ** indicates $p<0.01$ (ranksum test).

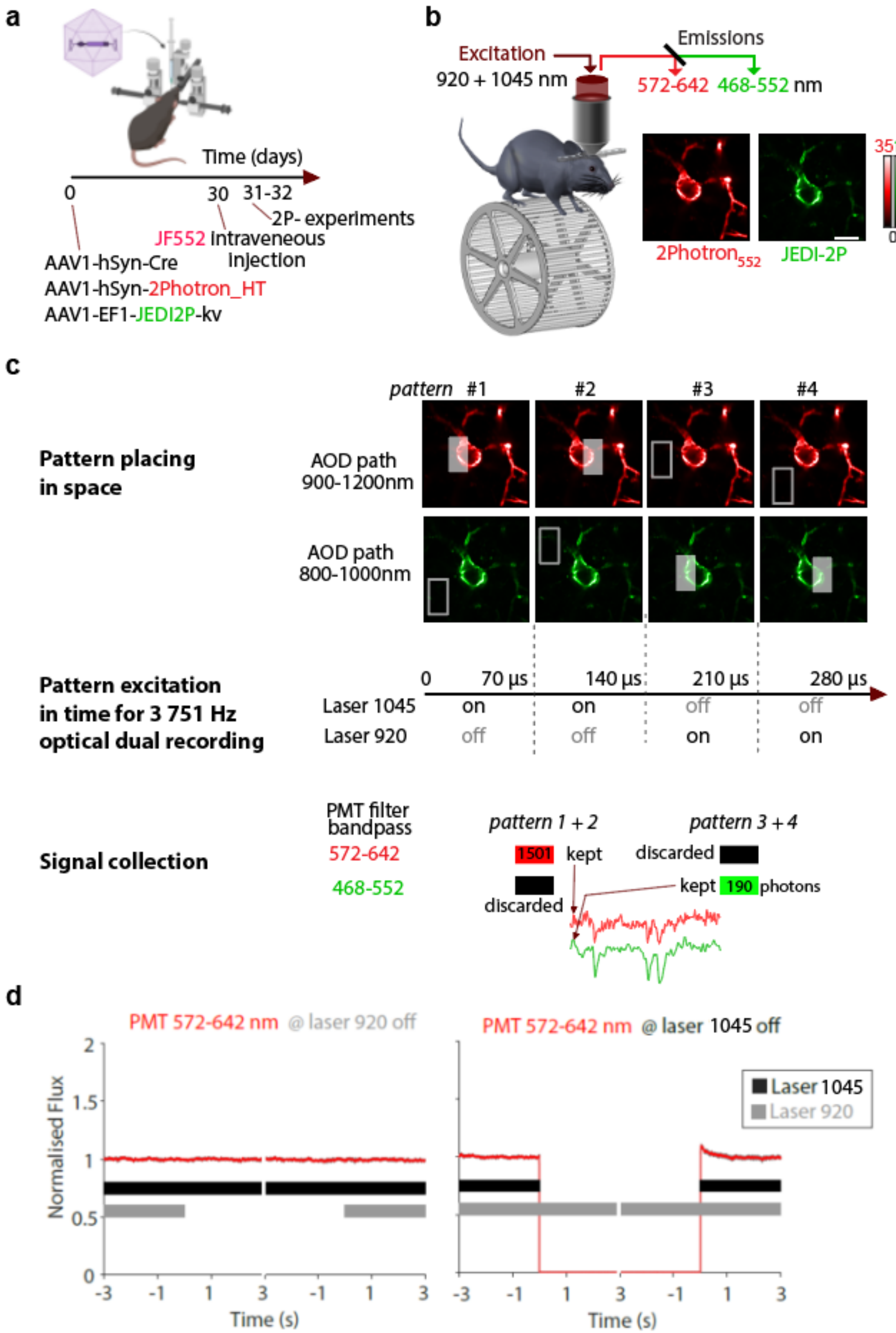
a
Time (days)
0
30
31-32
2P- experiments
JF552 intraveneous injection
AAV1-hSyn-Cre
AAV1-hSyn-2Photron_HT
AAV1-EF1-JEDI2P-kv
b
Excitation
920 + 1045 nm
Emissions
572-642 468-552 nm
3510
Photons
0
2Photron552
JEDI-2P
c
pattern #1 #2 #3 #4
Pattern placing in space
AOD path 900-1200nm
AOD path 800-1000nm
Pattern excitation in time for 3 751 Hz optical dual recording
0 70 µs 140 µs 210 µs 280 µs
Laser 1045 on on off off
Laser 920 off off on on
Signal collection
PMT filter bandpass
572-642
468-552
pattern 1 + 2
pattern 3 + 4
1501 kept
discarded
discarded
kept 190 photons
d
PMT 572-642 nm @ laser 920 off
PMT 572-642 nm @ laser 1045 off
Normalised Flux
Time (s)
Laser 1045
Laser 920

**Supplementary Figure 10. Method for multispectral two-photon optical recordings without spectral crosstalk. a**, Timeline of experimentation: virus injection preceded 2P-experiments by 30-40 days while JF552 fluorophore is injected 1 or 2 days before performing optical recordings. **b**, Left, experimental scheme and filter setting for cell recording. Right, single 2P-image planes of expression *in vivo*. **c**, Illustration of ULoVE excitation pattern positioning in space (top) and time (middle) used to obtain optical voltage records (bottom): Top row indicates how the four ULoVE patterns (70µs per pattern) are spatially placed for a single acquisition, two ULoVE patterns per cell are first applied through the 900-1200 nm AOD path then two other ULoVE patterns are applied through the 800-1000 nm path. To avoid cross-contamination of the signal pathways, patterns are alternately spatially offset and laser excitation is off. **d**, Normalized photon flux (n=14 cells) showing the absence of bleed-through from the 920 nm excitation into the 572-642 nm PMT, controlling for cross-contamination signals.

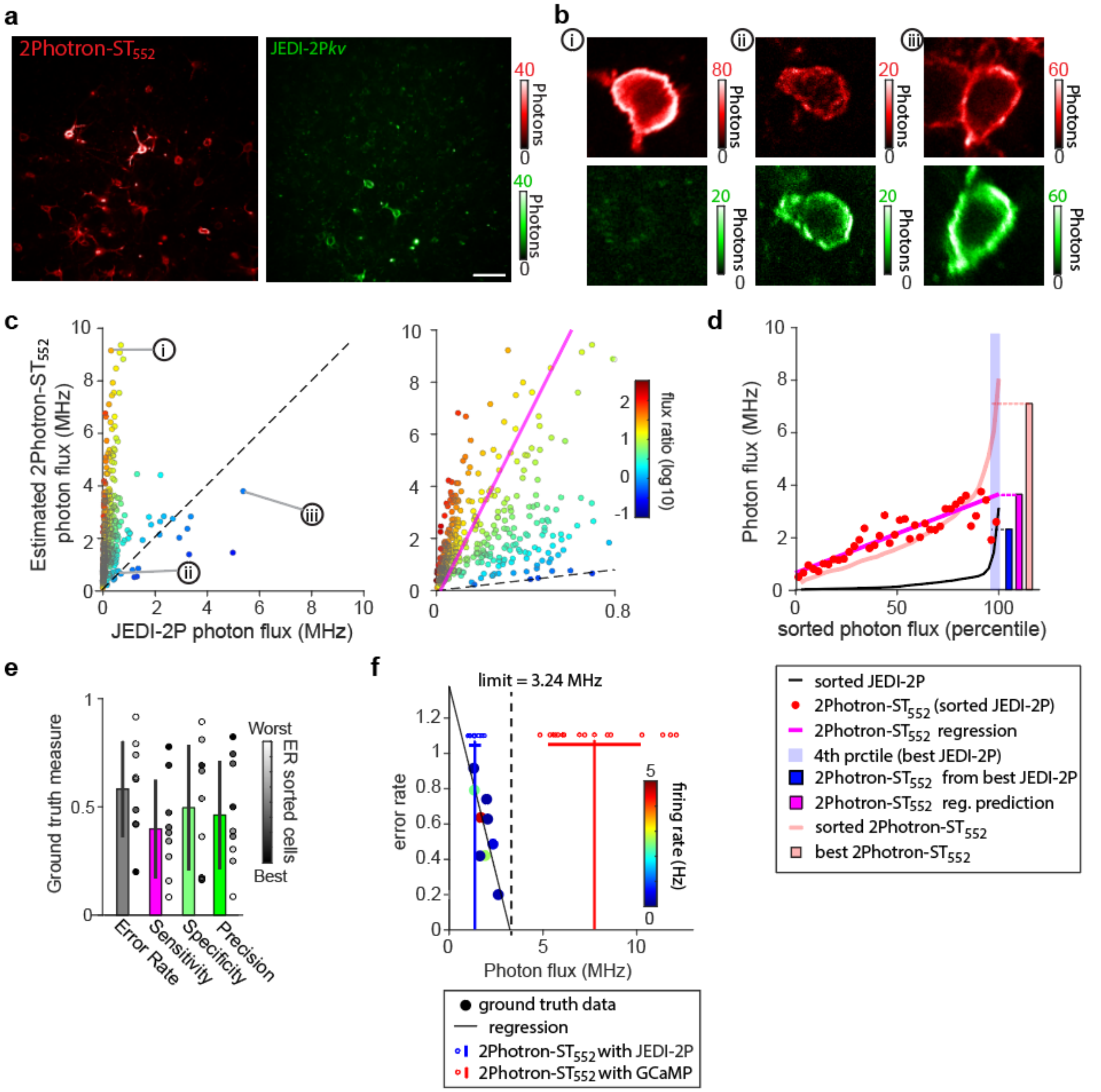


**Supplementary Figure 11. GEVI co-expression and ground truth calibration to determine ideal photon flux. a**, Full frame single-plane 2P images *in vivo* of 2Photron-$ST_{552}$ (left) and JEDI-2P-Kv (right), scale bar is 50 μm and depth is 214 μm below the meninges. **b**, Zoomed images for brightness levels of the different GEVIs, note differences of color scales. i,ii, and iii are examples displayed in the panel c, respectively high 2Photron-$ST_{552}$ with very low JEDI-2P, low 2Photron-$ST_{552}$ and low JEDI-2P, and average brightness levels for 2Photron-$ST_{552}$ and high JEDI-2P. **c**, Estimated 2Photron-$ST_{552}$ and JEDI-2P photon fluxes (n=403 neurons, 4 mice, depth: 165.62 ± 70.4 µm). Colors represent the log10 of flux ratio. Dashed line log10 flux ratio of 0. Note neurons expressing workable JEDI-2-Kv photon flux display only a weak 2Photron-$ST_{552}$ photon flux. Right, zoom where the magenta line represents the principal component of high ratio data points (above 5 folds). **d**, Mean photon flux (JEDI-2-Kv black, 2Photron-$ST_{552}$ red dots) as a function of JEDI-2-Kv flux expressed in percentile. Interestingly, we found: i) a correlation between JEDI2P-Kv and 2Photron-$ST_{552}$ photon fluxes. This correlation holds for the 96.3 first percent of the distribution (388/403 neurons, Pearson correlation coefficient: r= 0.92422, p-value 1.2623e-16, slope (magenta): 0.304 MHz / 10 % of ranked cell), ii) within the 3.7 last percent of the distribution when the JEDI-2P-Kv reached its highest flux, we found a sudden drop of the 2Photron-$ST_{552}$ photon flux (15/403 neurons, blue shaded area). When compared to the value predicted by the regression (thin magenta line), this drop corresponds to a flux decrease of 38.46% (predicted photon flux: 3.7 MHz, measured photon flux: 2.28 MHz). Note: for reasons still unclear, however, the highest expression of either indicator was not observed in cells of co-expression. Thus, cells of the highest JEDI2P-Kv expression were chosen to establish the ground truth spiking activity, even though this meant they are amongst the lower level of 2Photron co-expression and consequently limited the epochs of comparison to the first minute of the 10-minute recording. **e**, Measurements of Error rate. Sensitivity, Specificity, and Precision from spikes during the first minute of the JEDI-2P-Kv and 2Photron-$ST_{552}$ traces (n=9 cells, 4 mice). The error rate corresponds to the harmonic average of the sensitivity and the precision. Note that error rate values (sorted by gray shading) correlate with the overall metrics. **f**, Limit of the photon flux that enables a good theoretical detection. Plot of error rate as a function of average photon flux during the first minute of recordings. Note: regardless of cell firing rate, a lower error rate correlates with a higher photon flux. Linear regression projected to y=0 gives a limit of 3.24 MHz. Note that this limit of theoretically perfect detection is well below the distribution of the ULoVE-based 2Photron-$ST_{552}$ photon fluxes extracted from the 2Photron-$ST_{552}$ / GCaMP experiments (n=16 neurons).

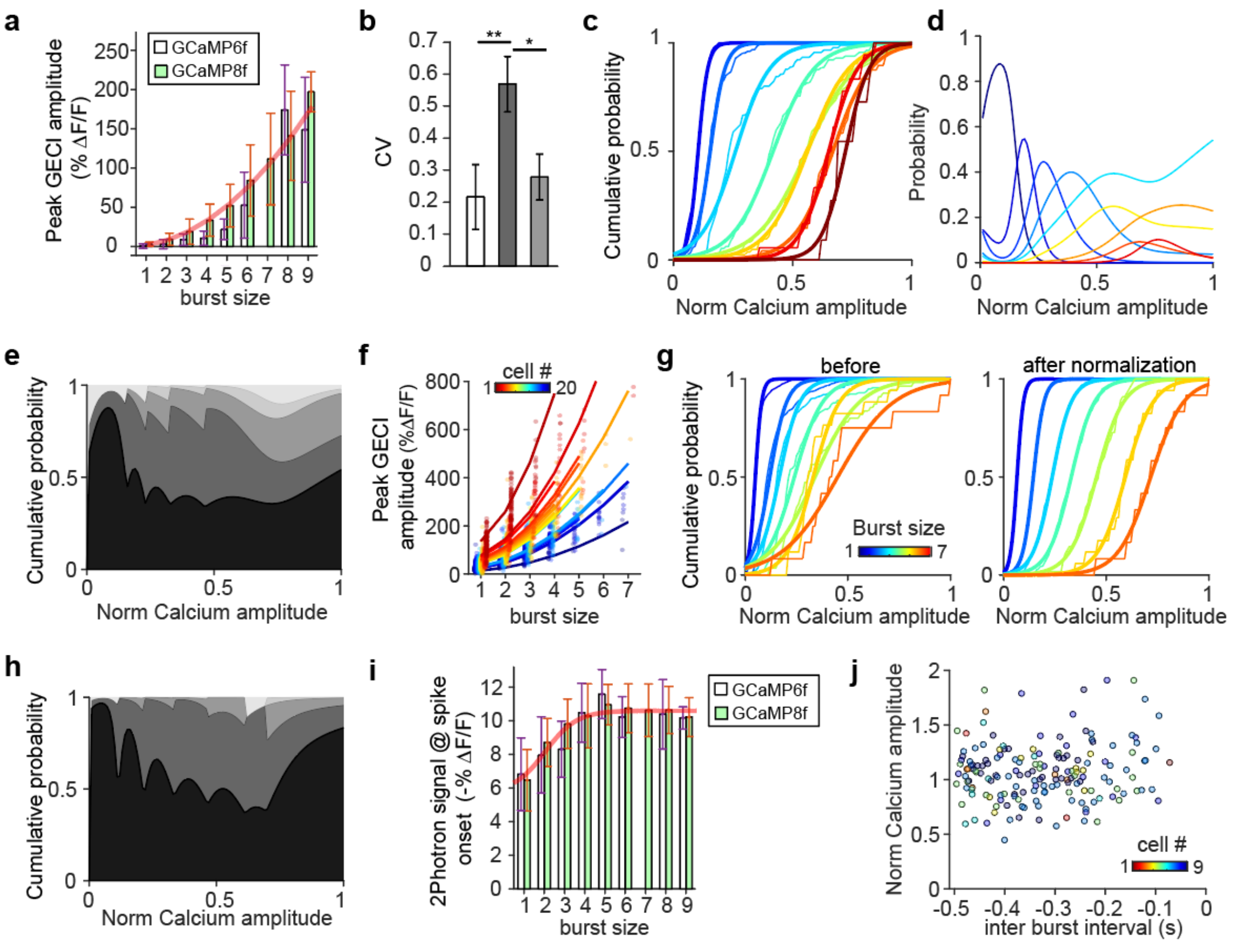


**Supplementary Figure 12. Normalizations and distributions applied to data of Zhang et al., 2023. a**, Calcium transient peak amplitude distribution versus burst size. White, GCaMP6f, green, GCaMP8f. Red, quadratic fit for GCaMP8f data. **b**, Mean burst size coefficient of variation. White, individual cells, Dark grey, cells pooled together, Light grey cells normalized and pooled). Rank sum test: ** p value <0.01, * pvalue < 0.05. error +/- std. **c** and **d**, As for Fig. 4**h** normalizing on the largest event of the cell without considering the burst size distribution. **e**, As for Fig. 4**J** using the distribution in **d**. **f**, Zhang et al., 2023 calcium transient amplitudes as a function of burst size for 20 individual GCaMP8f cells. Color code is according to a scaling factor obtained from quadratic fit, and ranged from 14 to 136 (mean +/- std: 46.4 +/- 26.5). **g**, As for Fig. 4**h**, using data taken from Zhang et al., 2023. Left, before normalization (CV: 0.569). Right, after normalization (CV: 0.279). **h**, As for Fig. 4, using data taken from Zhang et al., 2023. Despite fewer burst sizes, comparable results are obtained for true positive and average spike error per event: 68.9% and 0.291 (before normalization), 77.8% and 0.214 (after normalization), and 73% and 0.292 (for the normalization performed without knowing the burst size). **i**, Distribution of the associated 2Photron-$ST_{552}$ signals preceding the last spike of the burst. Red, sigmoid fit indicates that the spikes saturated after 5.01 spikes. **j**, Normalized calcium transient peak amplitude as a function of the interburst interval in the 2Photron-GCaMP8f dataset. Datapoints are color coded according to the neuron number from figure 4**g**.

**Table 1. Rhodopsin domain sequences used for fusion with HaloTag based on sequence alignment with the 7 transmembrane domain of Ace2 rhodopsin**

| Organism (abbreviation) | sequence |
|---|---|
| *Acetabularia acetabulum*<br><br>(Ace2) | MADVETETGMIAQWIVFAIMAAAAIAFGVAVHFRPSELKSAYYINIAICTIAATAYYAMA<br>VNYQDLTMNGERQVVYARYIDWVLTTPLLLLDLIVMTKMGGVMISWVIGADIFMIVFGIL<br>GAFEDEHKFKWVYFIAGCVMQAVLTYGMYNATWKDDLKKSPEYHSSYVSLLVFLSILWVF<br>YPVVWAFGSGSGVLSVDNEAILMGILDVLAKPLFGMGCLIAHETIFK |
| *Haloarcula argentinensis*<br>(Cruxrhodopsin-1)<br>(CR1) | MPEPGSEAIWLWLGTAGMFLGMLYFIARGWGETDSRRQKFYIATILITAIAFVNYLAMAL<br>GFGLTIVEFAGEEHPIYWARYSDWLFTTPLLLYDLGLLAGADRNTITSLVSLDVLMIGTG<br>LVATLSPGSGVLSAGAERLVWWGISTAFLLVLLYFLFSSLSGRVADLPSDTRSTFKTLRN<br>LVTVVWLVYPVWWLIGTEGIGLVGIGIETAGFMVIDLTAKVGFGIILLRSHGVLDGAA |
| *Leptosphaeria maculans*<br><br>(LmR) | MIVDQFEEVLMKTSQLFPLPTATQSAQPTHVAPVPTVLPDTPIYETVGDSGSKTLWVVFV<br>LMLIASAAFTALSWKIPVNRRLYHVITTIITLTAALSYFAMATGHGVALNKIVIRTQHDH<br>VPDTYETVYRQVYYARYIDWAITTPLLLLDLGLLAGMSGAHIFMAIVADLIMVLTGLFAA<br>FGSEGTPQKWGWYTIACIAYIFVVWHLVLNGGANARVKGEKLRSFFVAIGAYTLILWTAY<br>PIVWGLADGARKIGVDGEIIAYAVLDVLAKGVFGAWLLVTHANLRESD |
| *Podospora anserina*<br><br>(PaR) | MIHHDQVAEMLYGYAGAAAAKPTHTDSPGPIPTVIPTPPQFQEIGETGHRTLWVVFALMV<br>LSSGFFAFMSWNVPISKRLYHVITTLITITASLSYFAMASGHVTSFSCTPAKDHHKHVPD<br>VGYTECRQVFWGRYVDWAITTPLLLLDLSLLAGIDGAHTLMAVIADVIMVLSGLFASQGE<br>TATQRWGWYAIGCVSYLFVIWHVALHGARTVTAKGRGVTRLFSSLALFTFVLWTAYPIVW<br>GIADGAHRTTVDTEILIYAVLDILAKPVFGLWLLFSHRSLAETN |
| *Neurospora crassa*<br><br>(NcR) | MIHPEQVADMLRPTTSTTSSHVPGPVPTVVPTPTEYQTLGETGHRTLWVTFALMVLSSGI<br>FALLSWNVPTSKRLFHVITTLITVVASLSYFAMATGHATTFNCDTAWDHHKHVPDTSHQV<br>CRQVFWGRYVDWALTTPLLLLELCLLAGVDGAHTLMAIVADVIMVLCGLFAALGEGGNTA<br>QKWGWYTIGCFSYLFVIWHVALHGSRTVTAKGRGVSRLFTGLAVFALLLWTAYPIIWGIA<br>GGARRTNVDTEILIYTVLDLLAKPVFGFWLLLSHRAMPETN |
| *Acetabularia acetabulum*<br><br>(Ace1) | MSNPNPFQTTLGTDAQWVVFAVMALAAIVFSIAVQFRPLPLRLTYYVNIAICTIAATAYY<br>AMAVNGGDNKPTAGTGADERQVIYARYIDWVFTTPLLLLDLVLLTNMPATMIAWIMGADI<br>AMIAFGIIGAFTVGSYKWFYFVVGCIMLAVLAWGMINPIFKEELQKHKEYTGAYTTLLIY<br>LIVLWVIYPIVWGLGAGGHIIGVDVEIIAMGVLDLLAKPLYAIGVLITVEVVYGK |
| *Chlorella vulgaris*<br><br>(CvR) | MAVHQIGEGGLVMYWVTFGLMAFSALAFAVMTFTRPLNKRSHGYITLAIVTIAAIAYYAM<br>AASGGKALVSNPDGNLRDIYYARYIDWFFTTPLLLLDIILLTGIPIGVTLWIVLADVAMI<br>MLGLFGALSTNSYRWGYYGVSCAFFFVVLWGLFFPGAKGARARGGQVPGLYFGLAGYLAL<br>LWFGYPIVWGLAEGSDYISVTAEAASYAGLDIAAKVVFGWAVMLSHPLIARNQ |
| *Klebsormidium nitens*<br><br>(KnR) | MVYSHADAAGWTLYWITYGIMAVTALIFFAMSLRRPIQQRSHHYTSFLIVAIASLAYYAM<br>ASQGGNTRIRVYQGPADSYRQIFWARYVDWFFTTPLLLLDLVLLSNLSKLRIAAIMVADI<br>FMILTGLFGAVEARSNKWGWFVFGCIFMLYIFYELLVNVRKGAYARGGQHGMLYSVLLVW<br>LLILWVQYPVVWGLAEGSSTVSSDTEIAWYAALDICAKCVFGFILLLGIESIDRKR |
| *Taphrina deformans*<br><br>(TdR) | MSSLFEKRNTAVATNYRGPTNSIVINEAGSDWYWAVFSVMAASAIVFSVMAAMTPRGERV<br>FHYLTIAIVSVASVAYFTMAADLGSVAIISEFANYSSLPTRQVFYARYIDWVITTPLLLT<br>DLMLLAGLPWSTIIFTIVMDEVMVLTGLFGAITPSSYKWGYFTFGMVAYFFVAWVLIVEA<br>RKNAHRLGSDVHRLYIGIAIWTATLWTLYPVAWGLSEGGNVTSSDGEAIFYGVLDLLAKP<br>VFGLWILLGHKGIGMDRL |
| *Exophiala sideris*<br><br>(EsR) | MESNLVRRNGALTVNTMTQNNQSVAIHQTVRGSDWYYTVCAVMGTTSLAILALSRMKPRT<br>DRVFFYLTSGLCMVACIAYFAMGSNLGWTPIDVEWLRNDSVVRGVNRQVFYARYIDWVIT<br>TPMLLMDLLLTAGMPWPTILWIILLDEIMIVTGLIGALVKSRYKWGFYVFGCMAMFYIMW<br>ELAFPARKHAKVLGKDIHRSFVLCGVLTLVVWLCYPICWGLSEGGNVISPDSESVFYGVL<br>DVLAKPSFSIALIATHWNIDPGR |
| *Moelleriella libera*<br><br>(MlR) | MGNSAIEVNGDTVDSYTADVKITTHGSDAYWAITAVMAFTTILFIAHSFTKPRTDRIFHY<br>ITISITLVASIAYFTMASNLGWASIFIEFQRDDPLVSGTTREIFYVRYIDWVITTPLLLL<br>DILLTAGLPWPTILFTILLDEVMIITGLVGALVKSSYKWGFFTFGCVAFFGVAWSVAWTG<br>RKHANALGSDIGRVYLMTSVWTLFLWLLYPIAWGVSEGGNVISPDSEAAFYGTLDVLAKP<br>IFGIILLWGHRNIPASR |
| *Saitoella complicate*<br><br>(ScR) | MDSLLLQRRNDAVATNPPNATFALTENGSSWLWAVFCVMALSCIIIAALSLFKPMGYRIF<br>YLLNVAILATASVSYFSLASDLGLTPVTVEFRGPGTRQIAYVRYIDWFVTTPLLLTELLL<br>TAGLPTNIIISTIFADLVMIITGLAGALVVSRYKWGYYTMGCVAMLWVFWNVFTGIKVSG<br>NIGPDVRKSYTLLAVWLMIIWLNYPICWGLAEGGNRITVVGEMVYYGVLDLLAKPVFAAI<br>SLAVHSKIELSR |

| | |
|---|---|
| *Rhodotorula graminis*<br><br>(RgR) | MDAILSKRNEVLSLNPLVANIDITTAASDWLWAVFAVMGLSAIILLVLGHATRPIGERAF<br>HELAAALCFTASIAYYSMASDLGATPIEVEFIRGGTLGQNWVDIGVLRPTRSIWYARYID<br>WTITTPLLLLELALTTALPLSQIFGLVFFDIVMIITGLLGALTASRYKWGFFVFGCVAMF<br>WIFWVLFFPARKSASHLGTDYHRAYTSSAIVLCTLWTVYPIIWGVCDGGNVITPTSEMVA<br>YGVLDLLAKPVFSFWHVFQLSRLDYAR |
| *Rhodotorula toruloides*<br><br>(RtR) | MPFYIKNADIDITTHGSDWLWAVFSVMLLSAIGILVWGHVARPLGERAFHELAAALCFTA<br>SIAYFAMASDLGDVPIVVEFIRGGSLGQNWVQVGVENPTRAIWYARYIDWTITTPMLLLE<br>LLLCTGLPLSQVFSVIFADLLMIETGLIGALVASRYKWGFYAFGCAAQLYIWWMLLVPGR<br>RSAQHIGSDFAKSYTMSNIFLTTVWLVYPVIWGVADGGNVITPDSEMIAYGVLDLLAKPV<br>FSVIHLMSLSKLDYAR |
| *Krokinobacter eikastus*<br><br>(KeR1) | MKFLLLLLADPTKLDPSDYVGFTFFVGAMAMMAASAFFFLSLNQFNKKWRTSVLVSGLIT<br>FIAAVHYWYMRDYWFAIQESPTFFRYVDWVLTVPLMCVEFYLILKVAGAKPALMWKLILF<br>SVIMLVTGYFGEAVFQDQAALWGAISGAAYFYIVYEIWLGSAKKLAVAAGGDILKAHKIL<br>CWFVLVGWAIYPLGYMLGTDGWYTSILGKGSVDVAYNIADAINKIGFGLVIYALAVKKNE<br>VD |
| *Krokinobacter eikastus*<br><br>(KeR2) | MTQELGNANFENFIGATEGFSEIAYQFTSHILTLGYAVMLAGLLYFILTIKNVDKKFQMS<br>NILSAVVMVSAFLLLYAQAQNWTSSFTFNEEVGRYFLDPSGDLFNNGYRYLNWLIDVPML<br>LFQILFVVSLTTSKFSSVRNQFWFSGAMMIITGYIGQFYEVSNLTAFLVWGAISSAFFFH<br>ILWVMKKVINEGKEGISPAGQKILSNIWILFLISWTLYPGAYLMPYLTGVDGFLYSEDGV<br>MARQLVYTIADVSSKVIYGVLLGNLAITLSKNK |
| *Gloeobacter violaceus*<br><br>(GvR) | MLMTVFSSAPELALLGSTFAQVDPSNLSVSDSLTYGQFNLVYNAFSFAIAAMFASALFFF<br>SAQALVGQRYRLALLVSAIVVSIAGYHYFRIFNSWDAAYVLENGVYSLTSEKFNDAYRYV<br>DWLLTVPLLLVETVAVLTLPAKEARPLLIKLTVASVLMIATGYPGEISDDITTRIIWGTV<br>STIPFAYILYVLWVELSRSLVRQPAAVQTLVRNMRWLLLLSWGVYPIAYLLPMLGVSGTS<br>AAVGVQVGYTIADVLAKPVFGLLVFAIALVKTKAD |
| *Salinibacter ruber*<br><br>(SrR) | MLQELPTLTPGQYSLVFNMFSFTVATMTASFVFFVLARNNVAPKYRISMMVSALVVFIAG<br>YHYFRITSSWEAAYALQNGMYQPTGELFNDAYRYVDWLLTVPLLTVELVLVMGLPKNERG<br>PLAAKLGFLAALMIVLGYPGEVSENAALFGTRGLWGFLSTIPFVWILYILFTQLGDTIQR<br>QSSRVSTLLGNARLLLLATWGFYPIAYMIPMAFPEAFPSNTPGTIVALQVGYTIADVLAK<br>AGYGVLIYNIAKAKSEEE |
| *Gammaproteobacteria*<br><br>(GpR1) | MKLLLILGSVIALPTFAAGGGDLDASDYTGVSFWLVTAALLASTVFFFVERDRVSAKWKT<br>SLTVSGLVTGIAFWHYMYMRGVWIETGDSPTVFRYIDWLLTVPLLICEFYLILAAATNVA<br>GSLFKKLLVGSLVMLVFGYMGEAGIMAAWPAFIIGCLAWVYMIYELWAGEGKSACNTASP<br>AVQSAYNTMMYIIIFGWAIYPVGYFTGYLMGDGGSALNLNLIYNLADFVNKILFGLIIWN<br>VAVKESSN |
| *Gammaproteobacteria*<br><br>(GpR2) | MGKLLLILGSAIALPSFAAAGGDLDISDTVGVSFWLVTAGMLAATVFFFVERDQVSAKWK<br>TSLTVSGLITGIAFWHYLYMRGVWIDTGDTPTVFRYIDWLLTVPLQVVEFYLILAACTSV<br>AASLFKKLLAGSLVMLGAGFAGEAGLAPVLPAFIIGMAGWLYMIYELYMGEGKAAVSTAS<br>PAVNSAYNAMMMIIVVGWAIYPAGYAAGYLMGGEGVYASNLNLIYNLADFVNKILFGLII<br>WNVAVKESSN |
| *Proteomonas sulcata*<br><br>(PsuCCR) | MTMLEHLEGTMDGWYAENDLGQGAIIAHWVTFFFHMITTFYLGYVSFHSKGPGGKQPYFA<br>GYHEENNIGIFVNLFAAISYFGKVVSDTHGHNYQNVGPFIIGLGNYRYADYMLTCPLLVM<br>NLLFQLRAPYKITCAMLIFAVLMIGAVTNFYPGDDMKGPAVAWFCFGCFWYLIAYIFMAH<br>IVSKQYGRLDYLAHGTKAEGALFSLKLAIITFFAIWVAFPLVWLLSVGTGVLSNEAAEIC<br>HCICDVVAKSVYGFALANFREQYDREL |
| Guillardia theta<br><br>(GtCCR1) | MVESSAVIAANWISFLVIAGSFVVLCFISLRYKGPGGNENYYNGFREQNMLTVIINLWCA<br>LAYFAKVLQSHSDDDGFVPLTKIPYLDYATTCPLLTLDLMWCLDAPYKITSAVLVFTVMI<br>TGVACSLAVAPYSFYWFAMGMVLFIFTYVLMLSIVRERLEFITQCAHDSNAKRSIKHLKA<br>AVIIYFGIWPIFAILWLLSYRAANVISNDTNHILHCILDVIAKSCFGFVLLHFKMYFDKK<br>L |
| Guillardia theta<br><br>(GtCCR2) | MVASSAVITANWISFLAISASFIILLVISLRYKGPGGTESFYNGFKEQNMLTVFINLWCA<br>LAYFAKVLQSHSNDNGFAPLTVIPYVDYCTTCPLLTLDLLWCLDAPYKISSAVLVFTCLV<br>IAVACSLAVAPFSYCWFAMGMVLFTFTYVFILSIVRQRLDFFTLCARDSNAKQSLKHLKT<br>AVFIYFGIWLLFPLLWLLSYRAANVISNDINHIFHCILDVIAKSVYGFALLYFKMYFDKK<br>L |
| Guillardia theta<br><br>(GtCCR3) | MVSALDQNGPQYLQNPIVIAADWIGFIALFGSSLAVAYKLVTFKGPDQDDVYFFGYREEK<br>MISVFVNLFAALAYWAKLASHANGDVGPAASVTTYKYLDYLFTCPLLTIDLLWCLNLPYK<br>FTFGAIVAVCILCAFMASVIPPPARYMWFGMGITVFSAAWFNILKLVRMRLEQFVSKEAK<br>KVRQSLKVACMTYFFIWLGYPTLWVLGDAGVLDSVVSALLHTFLDVFSKSIYGFALLHFV<br>MRTDKRE |
| Guillardia theta | MTTSAPSLSDPNWQYGMGGWNNPRLPNFNLHDPTVIGVDWLGFLCLLGASLALMYKLMSF<br>KGPDGDQEFFVGYREEKCLSIYVNLIAAITYWGRICAHFNNDMGLSLSVNYFKYLDYIFT |

| | |
|---|---|
| (GtCCR4) | CPILTLDLLWSLNLPYKITYSLFVGLTIACNAFEPPARYLWFMFGCFIFAFTWISIIRLVYARFQQFLNEDAKKIRAPLKLSLTLYFSIWCGYPALWLLTEFGAISQLAAHVMTVIMDVAAKSVYGFALLKFQLGVDKRD |
| Guillardia theta<br><br>(GtCCR5) | MSTSSVAYLRTPVVQALDWVGFISLGGTAAVLAYRLMNFKPPNKDILYFFGYREKGMISLYVNLFAAVAYYARITSHLSGDVGAATNIILYKYFDYLITCPLLTFDLLTTLNLPYKITYAVYVQITIFTGFMSANTPPPATFLWFAFGMLLFSYTWFNIISLVQVRFIQYFAKKGNTTQSRRVSVASKAGFRNKNVRNPLQTALSTYFCIWMVYPVLWLLLKTKVIDQVTEHCINVVMDVLAKSMYGFALLRFQLLMDKAN |
| *Emiliania huxleyi*<br><br>(EhR) | MDTIVLTINWFNFIAGLFHVALAIVCALLGDVSQTFKMYMPFTTFRNGTQENEVTIKYSGYFPMTVIFVVYFSVTALFHMGNAFLWNDTYHRFLSNRKNPIRWTEYSITAPLMTAILAFIAGSRNWLFVIAASTLTFASISVGFVLDEIRKYSFFVLGMIPFLVEFSLIILSLYLTTSCYPSYLPITIFVEFGLWILFPFVSLLELSGINYKIGELVFIVLSFVSKSVLAIILNSENVLKNGVTGIC |
| *Actinomycetes bacterium*<br><br>(AbR) | MAKPTVKEIKSLQNFNRIAGVFHLLQMLAVLALANDFALPMTGTYLNGPPGTTFSAPVVILETPVGLAVALFLGLSALFHFIVSSGNFFKRYSASLMKNQNIFRWVEYSLSSSVMIVLIAQICGIADIVALLAIFGVNASMILFGWLQEKYTQPKDGDLLPFWFGCIAGIVPWIGLLIYVIAPGSTSDVAVPGFVYGIIISLFLFFNSFALVQYLQYKGKGKWSNYLRGERAYIVLSLVAKSALAWQIFSGTLIP |
| *Heimdallarchaeota archaeon*<br><br>(HaR) | MTKALEMTILDDGSESPISFKYLRKFNLAMGILHLVQGVAMLVLGFFWDFSRPLYSITLDYSGGPPVAALQPEFSFTAVGPTVAAFLLFSALAHLLIAGPLNKFYVKNLKKKMNPIRWFEYAFSSSIMVFFIAILFGVWDLWVLIGLFFLNMLMNLFGHMMELHNQTTKKTNWTAYIYGWIAGIIPWVIISVFFARIAINSTGMPWFVPVIYAFELILFMSFAFNMLLQYKKVGKWKDYLYGERMYQILSLVAKTLLAWLVFAGVFQPA |
| *Halobacterium sp.DL1*<br><br>(HdR) | MSSHSATTSSGRRESARDSRLRLWNSVMAVLHFLQGAAMVLLADTVLWPITRTRYGFDPGSQSIFPETVAFVDANLPLLVAGFLFISALAHTAIATVWYDKYVRYLDRGMNPYRWYEYSVSASLMIVVIGMLSGVWDLGTLVALFGLVAVMNLSGLLMEQRNELTEQTDWTPYWVGVIAGIVPWITIGVAFVGSVTASAGEFPEFVIYIYVSIFVFFNLFALNMALQYLEVSRWKNYLFGEKMYIVLSLVAKSALAWQVYFGTLNSPI |
| *Thermoplasmatales archaeon SG8*<br><br>(TsR) | MTDYEDHTIRRFRIFNAIMGGIHLLQVFLVLYLSNNFSLPVTISKPVYNEFTNSISPVSETLFSIRVGPLVALFLFISAIAHILIATVLYYRYVENLKSGMNPYRWFEYSISASVMIVIIAMLTTIYDLGTLLALFTLTAVMNLMGLMMELHNQTTQNTDWTSYIIGCIAGLVPWIVIFIPLIAAESVPDFVIYIFVSIAIFFNCFAINMYLQYKKIGKWKDYLHGERVYIILSLVAKSALAWQVFAGTLRPM |

**Table 2: Protein sequence of all constructs generated by fusing HaloTag to the c-terminus of rhodopsins tested and introducing mutations to generate putative negative-going or positive-going voltage indicators**

| Construct Name | Sequence |
|---|---|
| CR1 (-) | MPEPGSEAIWLWLGTAGMFLGMLYFIARGWGETDSRRQKFYIATILITAIAFVNYLAMAL<br>GFGLTIVEFAGEEHPIYWARYSNWLFTTPLLLYDLGLLAGADRNTITSLVSLDVLMIGTG<br>LVATLSPGSGVLSAGAERLVWWGISTAFLLVLLYFLFSSLSGRVADLPSDTRSTFKTLRN<br>LVTVVWLVYPVWWLIGTEGIGLVGIGIETAGFMVIDLTAKVGFGIILLRSHGVLDGAAIG<br>TGFPFDPHYVEVLGERMHYVDVGPRDGTPVLFLHGNPTSSYVWRNIIPHVAPTHRCIAPD<br>LIGMGKSDKPDLGYFFDDHVRFMDAFIEALGLEEVVLVIHDWGSALGFHWAKRNPERVKG<br>IAFMEFIRPIPTWDEWPEFARETFQAFRTTDVGRKLIIDQNVFIEGTLPMGVVRPLTEVE<br>MDHYREPFLNPVDREPLWRFPNELPIAGEPANIVALVEEYMDWLHQSPVPKLLFWGTPGV<br>LIPPAEAARLAKSLPNCKAVDIGPGLNLLQEDNPDLIGSEIARWLSTLEISGEPTTKSRI<br>TSEGEYIPLDQIDINVFCYENEV |
| LmR (-) | MIVDQFEEVLMKTSQLFPLPTATQSAQPTHVAPVPTVLPDTPIYETVGDSGSKTLWVVFV<br>LMLIASAAFTALSWKIPVNRRLYHVITTIITLTAALSYFAMATGHGVALNKIVIRTQHDH<br>VPDTYETVYRQVYYARYINWAITTPLLLLDLGLLAGMSGAHIFMAIVADLIMVLTGLFAA<br>FGSEGTPQKWGWYTIACIAYIFVVWHLVLNGGANARVKGEKLRSFFVAIGAYTLILWTAY<br>PIVWGLADGARKIGVDGEIIAYAVLDVLAKGVFGAWLLVTHANLRESDIGTGFPFDPHYV<br>EVLGERMHYVDVGPRDGTPVLFLHGNPTSSYVWRNIIPHVAPTHRCIAPDLIGMGKSDKP<br>DLGYFFDDHVRFMDAFIEALGLEEVVLVIHDWGSALGFHWAKRNPERVKGIAFMEFIRPI<br>PTWDEWPEFARETFQAFRTTDVGRKLIIDQNVFIEGTLPMGVVRPLTEVEMDHYREPFLN<br>PVDREPLWRFPNELPIAGEPANIVALVEEYMDWLHQSPVPKLLFWGTPGVLIPPAEAARL<br>AKSLPNCKAVDIGPGLNLLQEDNPDLIGSEIARWLSTLEISGEPTTKSRITSEGEYIPLD<br>QIDINVFCYENEV |
| PaR (-) | MIHHDQVAEMLYGYAGAAAAKPTHTDSPGPIPTVIPTPPQFQEIGETGHRTLWVVFALMV<br>LSSGFFAFMSWNVPISKRLYHVITTLITITASLSYFAMASGHVTSFSCTPAKDHHKHVPD<br>VGYTECRQVFWGRYVNWAITTPLLLLDLSLLAGIDGAHTLMAVIADVIMVLSGLFASQGE<br>TATQRWGWYAIGCVSYLFVIWHVALHGARTVTAKGRGVTRLFSSLALFTFVLWTAYPIVW<br>GIADGAHRTTVDTEILIYAVLDILAKPVFGLWLLFSHRSLAETNIGTGFPFDPHYVEVLG<br>ERMHYVDVGPRDGTPVLFLHGNPTSSYVWRNIIPHVAPTHRCIAPDLIGMGKSDKPDLGY<br>FFDDHVRFMDAFIEALGLEEVVLVIHDWGSALGFHWAKRNPERVKGIAFMEFIRPIPTWD<br>EWPEFARETFQAFRTTDVGRKLIIDQNVFIEGTLPMGVVRPLTEVEMDHYREPFLNPVDR<br>EPLWRFPNELPIAGEPANIVALVEEYMDWLHQSPVPKLLFWGTPGVLIPPAEAARLAKSL<br>PNCKAVDIGPGLNLLQEDNPDLIGSEIARWLSTLEISGEPTTKSRITSEGEYIPLDQIDI<br>NVFCYENEV |
| NcR (-) | MIHPEQVADMLRPTTSTTSSHVPGPVPTVVPTPTEYQTLGETGHRTLWVTFALMVLSSGI<br>FALLSWNVPTSKRLFHVITTLITVVASLSYFAMATGHATTFNCDTAWDHHKHVPDTSHQV<br>CRQVFWGRYVNWALTTPLLLLELCLLAGVDGAHTLMAIVADVIMVLCGLFAALGEGGNTA<br>QKWGWYTIGCFSYLFVIWHVALHGSRTVTAKGRGVSRLFTGLAVFALLLWTAYPIIWGIA<br>GGARRTNVDTEILIYTVLDLLAKPVFGFWLLLSHRAMPETNIGTGFPFDPHYVEVLGERM<br>HYVDVGPRDGTPVLFLHGNPTSSYVWRNIIPHVAPTHRCIAPDLIGMGKSDKPDLGYFFD<br>DHVRFMDAFIEALGLEEVVLVIHDWGSALGFHWAKRNPERVKGIAFMEFIRPIPTWDEWP<br>EFARETFQAFRTTDVGRKLIIDQNVFIEGTLPMGVVRPLTEVEMDHYREPFLNPVDREPL<br>WRFPNELPIAGEPANIVALVEEYMDWLHQSPVPKLLFWGTPGVLIPPAEAARLAKSLPNC<br>KAVDIGPGLNLLQEDNPDLIGSEIARWLSTLEISGEPTTKSRITSEGEYIPLDQIDINVF<br>CYENEV |
| Ace1 (-) | MSNPNPFQTTLGTDAQWVVFAVMALAAIVFSIAVQFRPLPLRLTYYVNIAICTIAATAYY<br>AMAVNGGDNKPTAGTGADERQVIYARYINWVFTTPLLLLDLVLLTNMPATMIAWIMGADI<br>AMIAFGIIGAFTVGSYKWFYFVVGCIMLAVLAWGMINPIFKEELQKHKEYTGAYTTLLIY<br>LIVLWVIYPIVWGLGAGGHIIGVDVEIIAMGVLDLLAKPLYAIGVLITVEVVYGKIGTGF<br>PFDPHYVEVLGERMHYVDVGPRDGTPVLFLHGNPTSSYVWRNIIPHVAPTHRCIAPDLIG<br>MGKSDKPDLGYFFDDHVRFMDAFIEALGLEEVVLVIHDWGSALGFHWAKRNPERVKGIAF<br>MEFIRPIPTWDEWPEFARETFQAFRTTDVGRKLIIDQNVFIEGTLPMGVVRPLTEVEMDH<br>YREPFLNPVDREPLWRFPNELPIAGEPANIVALVEEYMDWLHQSPVPKLLFWGTPGVLIP<br>PAEAARLAKSLPNCKAVDIGPGLNLLQEDNPDLIGSEIARWLSTLEISGEPTTKSRITSE<br>GEYIPLDQIDINVFCYENEV |
| CvR (-) | MAVHQIGEGGLVMYWVTFGLMAFSALAFAVMTFTRPLNKRSHGYITLAIVTIAAIAYYAM |

| | |
|---|---|
| | AASGGKALVSNPDGNLRDIYYARYINWFFTTPLLLLDIILLTGIPIGVTLWIVLADVAMI<br>MLGLFGALSTNSYRWGYYGVSCAFFFVVLWGLFFPGAKGARARGGQVPGLYFGLAGYLAL<br>LWFGYPIVWGLAEGSDYISVTAEAASYAGLDIAAKVVFGWAVMLSHPLIARNQIGTGFPF<br>DPHYVEVLGERMHYVDVGPRDGTPVLFLHGNPTSSYVWRNIIPHVAPTHRCIAPDLIGMG<br>KSDKPDLGYFFDDHVRFMDAFIEALGLEEVVLVIHDWGSALGFHWAKRNPERVKGIAFME<br>FIRPIPTWDEWPEFARETFQAFRTTDVGRKLIIDQNVFIEGTLPMGVVRPLTEVEMDHYR<br>EPFLNPVDREPLWRFPNELPIAGEPANIVALVEEYMDWLHQSPVPKLLFWGTPGVLIPPA<br>EAARLAKSLPNCKAVDIGPGLNLLQEDNPDLIGSEIARWLSTLEISGEPTTKSRITSEGE<br>YIPLDQIDINVFCYENEV |
| KnR (-) | MVYSHADAAGWTLYWITYGIMAVTALIFFAMSLRRPIQQRSHHYTSFLIVAIASLAYYAM<br>ASQGGNTRIRVYQGPADSYRQIFWARYVNWFFTTPLLLLDLVLLSNLSKLRIAAIMVADI<br>FMILTGLFGAVEARSNKWGWFVFGCIFMLYIFYELLVNVRKGAYARGGQHGMLYSVLLVW<br>LLILWVQYPVVWGLAEGSSTVSSDTEIAWYAALDICAKCVFGFILLLGIESIDRKRIGTG<br>FPFDPHYVEVLGERMHYVDVGPRDGTPVLFLHGNPTSSYVWRNIIPHVAPTHRCIAPDLI<br>GMGKSDKPDLGYFFDDHVRFMDAFIEALGLEEVVLVIHDWGSALGFHWAKRNPERVKGIA<br>FMEFIRPIPTWDEWPEFARETFQAFRTTDVGRKLIIDQNVFIEGTLPMGVVRPLTEVEMD<br>HYREPFLNPVDREPLWRFPNELPIAGEPANIVALVEEYMDWLHQSPVPKLLFWGTPGVLI<br>PPAEAARLAKSLPNCKAVDIGPGLNLLQEDNPDLIGSEIARWLSTLEISGEPTTKSRITS<br>EGEYIPLDQIDINVFCYENEV |
| TdR (-) | MSSLFEKRNTAVATNYRGPTNSIVINEAGSDWYWAVFSVMAASAIVFSVMAAMTPRGERV<br>FHYLTIAIVSVASVAYFTMAADLGSVAIISEFANYSSLPTRQVFYARYINWVITTPLLLT<br>DLMLLAGLPWSTIIFTIVMDEVMVLTGLFGAITPSSYKWGYFTFGMVAYFFVAWVLIVEA<br>RKNAHRLGSDVHRLYIGIAIWTATLWTLYPVAWGLSEGGNVTSSDGEAIFYGVLDLLAKP<br>VFGLWILLGHKGIGMDRLIGTGFPFDPHYVEVLGERMHYVDVGPRDGTPVLFLHGNPTSS<br>YVWRNIIPHVAPTHRCIAPDLIGMGKSDKPDLGYFFDDHVRFMDAFIEALGLEEVVLVIH<br>DWGSALGFHWAKRNPERVKGIAFMEFIRPIPTWDEWPEFARETFQAFRTTDVGRKLIIDQ<br>NVFIEGTLPMGVVRPLTEVEMDHYREPFLNPVDREPLWRFPNELPIAGEPANIVALVEEY<br>MDWLHQSPVPKLLFWGTPGVLIPPAEAARLAKSLPNCKAVDIGPGLNLLQEDNPDLIGSE<br>IARWLSTLEISGEPTTKSRITSEGEYIPLDQIDINVFCYENEV |
| EsR (-) | MESNLVRRNGALTVNTMTQNNQSVAIHQTVRGSDWYYTVCAVMGTTSLAILALSRMKPRT<br>DRVFFYLTSGLCMVACIAYFAMGSNLGWTPIDVEWLRNDSVVRGVNRQVFYARYINWVIT<br>TPMLLMDLLLTAGMPWPTILWIILLDEIMIVTGLIGALVKSRYKWGFYVFGCMAMFYIMW<br>ELAFPARKHAKVLGKDIHRSFVLCGVLTLVVWLCYPICWGLSEGGNVISPDSESVFYGVL<br>DVLAKPSFSIALIATHWNIDPGRIGTGFPFDPHYVEVLGERMHYVDVGPRDGTPVLFLHG<br>NPTSSYVWRNIIPHVAPTHRCIAPDLIGMGKSDKPDLGYFFDDHVRFMDAFIEALGLEEV<br>VLVIHDWGSALGFHWAKRNPERVKGIAFMEFIRPIPTWDEWPEFARETFQAFRTTDVGRK<br>LIIDQNVFIEGTLPMGVVRPLTEVEMDHYREPFLNPVDREPLWRFPNELPIAGEPANIVA<br>LVEEYMDWLHQSPVPKLLFWGTPGVLIPPAEAARLAKSLPNCKAVDIGPGLNLLQEDNPD<br>LIGSEIARWLSTLEISGEPTTKSRITSEGEYIPLDQIDINVFCYENEV |
| MlR (-) | MGNSAIEVNGDTVDSYTADVKITTHGSDAYWAITAVMAFTTILFIAHSFTKPRTDRIFHY<br>ITISITLVASIAYFTMASNLGWASIFIEFQRDDPLVSGTTREIFYVRYINWVITTPLLLL<br>DILLTAGLPWPTILFTILLDEVMIITGLVGALVKSSYKWGFFTFGCVAFFGVAWSVAWTG<br>RKHANALGSDIGRVYLMTSVWTLFLWLLYPIAWGVSEGGNVISPDSEAAFYGTLDVLAKP<br>IFGIILLWGHRNIPASRIGTGFPFDPHYVEVLGERMHYVDVGPRDGTPVLFLHGNPTSSY<br>VWRNIIPHVAPTHRCIAPDLIGMGKSDKPDLGYFFDDHVRFMDAFIEALGLEEVVLVIHD<br>WGSALGFHWAKRNPERVKGIAFMEFIRPIPTWDEWPEFARETFQAFRTTDVGRKLIIDQN<br>VFIEGTLPMGVVRPLTEVEMDHYREPFLNPVDREPLWRFPNELPIAGEPANIVALVEEYM<br>DWLHQSPVPKLLFWGTPGVLIPPAEAARLAKSLPNCKAVDIGPGLNLLQEDNPDLIGSEI<br>ARWLSTLEISGEPTTKSRITSEGEYIPLDQIDINVFCYENEV |
| ScR (-) | MDSLLLQRRNDAVATNPPNATFALTENGSSWLWAVFCVMALSCIIIAALSLFKPMGYRIF<br>YLLNVAILATASVSYFSLASDLGLTPVTVEFRGPGTRQIAYVRYINWFVTTPLLLTELLL<br>TAGLPTNIIISTIFADLVMIITGLAGALVVSRYKWGYYTMGCVAMLWVFWNVFTGIKVSG<br>NIGPDVRKSYTLLAVWLMIIWLNYPICWGLAEGGNRITVVGEMVYYGVLDLLAKPVFAAI<br>SLAVHSKIELSRIGTGFPFDPHYVEVLGERMHYVDVGPRDGTPVLFLHGNPTSSYVWRNI<br>IPHVAPTHRCIAPDLIGMGKSDKPDLGYFFDDHVRFMDAFIEALGLEEVVLVIHDWGSAL<br>GFHWAKRNPERVKGIAFMEFIRPIPTWDEWPEFARETFQAFRTTDVGRKLIIDQNVFIEG<br>TLPMGVVRPLTEVEMDHYREPFLNPVDREPLWRFPNELPIAGEPANIVALVEEYMDWLHQ<br>SPVPKLLFWGTPGVLIPPAEAARLAKSLPNCKAVDIGPGLNLLQEDNPDLIGSEIARWLS<br>TLEISGEPTTKSRITSEGEYIPLDQIDINVFCYENEV |
| RgR (-) | MDAILSKRNEVLSLNPLVANIDITTAASDWLWAVFAVMGLSAIILLVLGHATRPIGERAF<br>HELAAAALCFTASIAYYSMASDLGATPIEVEFIRGGTLGQNWVDIGVLRPTRSIWYARYIN<br>WTITTPLLLLELALTTALPLSQIFGLVFFDIVMIITGLLGALTASRYKWGFFVFGCVAMF<br>WIFWVLFFPARKSASHLGTDYHRAYTSSAIVLCTLWTVYPIIWGVCDGGNVITPTSEMVA |

| | |
|---|---|
| | YGVLDLLAKPVFSFWHVFQLSRLDYARIGTGFPFDPHYVEVLGERMHYVDVGPRDGTPVL<br>FLHGNPTSSYVWRNIIPHVAPTHRCIAPDLIGMGKSDKPDLGYFFDDHVRFMDAFIEALG<br>LEEVVLVIHDWGSALGFHWAKRNPERVKGIAFMEFIRPIPTWDEWPEFARETFQAFRTTD<br>VGRKLIIDQNVFIEGTLPMGVVRPLTEVEMDHYREPFLNPVDREPLWRFPNELPIAGEPA<br>NIVALVEEYMDWLHQSPVPKLLFWGTPGVLIPPAEAARLAKSLPNCKAVDIGPGLNLLQE<br>DNPDLIGSEIARWLSTLEISGEPTTKSRITSEGEYIPLDQIDINVFCYENEV |
| RtR (-) | MPFYIKNADIDITTHGSDWLWAVFSVMLLSAIGILVWGHVARPLGERAFHELAAALCFTA<br>SIAYFAMASDLGDVPIVVEFIRGGSLGQNWVQVGVENPTRAIWYARYINWTITTPMLLLE<br>LLLCTGLPLSQVFSVIFADLLMIETGLIGALVASRYKWGFYAFGCAAQLYIWWMLLVPGR<br>RSAQHIGSDFAKSYTMSNIFLTTVWLVYPVIWGVADGGNVITPDSEMIAYGVLDLLAKPV<br>FSVIHLMSLSKLDYARIGTGFPFDPHYVEVLGERMHYVDVGPRDGTPVLFLHGNPTSSYV<br>WRNIIPHVAPTHRCIAPDLIGMGKSDKPDLGYFFDDHVRFMDAFIEALGLEEVVLVIHDW<br>GSALGFHWAKRNPERVKGIAFMEFIRPIPTWDEWPEFARETFQAFRTTDVGRKLIIDQNV<br>FIEGTLPMGVVRPLTEVEMDHYREPFLNPVDREPLWRFPNELPIAGEPANIVALVEEYMD<br>WLHQSPVPKLLFWGTPGVLIPPAEAARLAKSLPNCKAVDIGPGLNLLQEDNPDLIGSEIA<br>RWLSTLEISGEPTTKSRITSEGEYIPLDQIDINVFCYENEV |
| GpR1 (-) | MKLLLILGSVIALPTFAAGGGDLDASDYTGVSFWLVTAALLASTVFFFVERDRVSAKWKT<br>SLTVSGLVTGIAFWHYMYMRGVWIETGDSPTVFRYINWLLTVPLLICEFYLILAAATNVA<br>GSLFKKLLVGSLVMLVFGYMGEAGIMAAWPAFIIGCLAWVYMIYELWAGEGKSACNTASP<br>AVQSAYNTMMYIIIFGWAIYPVGYFTGYLMGDGGSALNLNLIYNLADFVNKILFGLIIWN<br>VAVKESSNIGTGFPFDPHYVEVLGERMHYVDVGPRDGTPVLFLHGNPTSSYVWRNIIPHV<br>APTHRCIAPDLIGMGKSDKPDLGYFFDDHVRFMDAFIEALGLEEVVLVIHDWGSALGFHW<br>AKRNPERVKGIAFMEFIRPIPTWDEWPEFARETFQAFRTTDVGRKLIIDQNVFIEGTLPM<br>GVVRPLTEVEMDHYREPFLNPVDREPLWRFPNELPIAGEPANIVALVEEYMDWLHQSPVP<br>KLLFWGTPGVLIPPAEAARLAKSLPNCKAVDIGPGLNLLQEDNPDLIGSEIARWLSTLEI<br>SGEPTTKSRITSEGEYIPLDQIDINVFCYENEV |
| GpR2 (-) | MGKLLLLILGSAIALPSFAAAGGDLDISDTVGVSFWLVTAGMLAATVFFFVERDQVSAKWK<br>TSLTVSGLITGIAFWHYLYMRGVWIDTGDTPTVFRYINWLLTVPLQVVEFYLILAACTSV<br>AASLFKKLLAGSLVMLGAGFAGEAGLAPVLPAFIIGMAGWLYMIYELYMGEGKAAVSTAS<br>PAVNSAYNAMMMIIVVGWAIYPAGYAAGYLMGGEGVYASNLNLIYNLADFVNKILFGLII<br>WNVAVKESSNIGTGFPFDPHYVEVLGERMHYVDVGPRDGTPVLFLHGNPTSSYVWRNIIP<br>HVAPTHRCIAPDLIGMGKSDKPDLGYFFDDHVRFMDAFIEALGLEEVVLVIHDWGSALGF<br>HWAKRNPERVKGIAFMEFIRPIPTWDEWPEFARETFQAFRTTDVGRKLIIDQNVFIEGTL<br>PMGVVRPLTEVEMDHYREPFLNPVDREPLWRFPNELPIAGEPANIVALVEEYMDWLHQSP<br>VPKLLFWGTPGVLIPPAEAARLAKSLPNCKAVDIGPGLNLLQEDNPDLIGSEIARWLSTL<br>EISGEPTTKSRITSEGEYIPLDQIDINVFCYENEV |
| KeR1 (-) | MKFLLLLLLADPTKLDPSDYVGFTFFVGAMAMMAASAFFFLSLNQFNKKWRTSVLVSGLIT<br>FIAAVHYWYMRDYWFAIQESPTFFRYVNWVLTVPLMCVEFYLILKVAGAKPALMWKLILF<br>SVIMLVTGYFGEAVFQDQAALWGAISGAAYFYIVYEIWLGSAKKLAVAAGGDILKAHKIL<br>CWFVLVGWAIYPLGYMLGTDGWYTSILGKGSVDVAYNIADAINKIGFGLVIYALAVKKNE<br>VDIGTGFPFDPHYVEVLGERMHYVDVGPRDGTPVLFLHGNPTSSYVWRNIIPHVAPTHRC<br>IAPDLIGMGKSDKPDLGYFFDDHVRFMDAFIEALGLEEVVLVIHDWGSALGFHWAKRNPE<br>RVKGIAFMEFIRPIPTWDEWPEFARETFQAFRTTDVGRKLIIDQNVFIEGTLPMGVVRPL<br>TEVEMDHYREPFLNPVDREPLWRFPNELPIAGEPANIVALVEEYMDWLHQSPVPKLLFWG<br>TPGVLIPPAEAARLAKSLPNCKAVDIGPGLNLLQEDNPDLIGSEIARWLSTLEISGEPTT<br>KSRITSEGEYIPLDQIDINVFCYENEV |
| KeR2 (-) | MTQELGNANFENFIGATEGFSEIAYQFTSHILTLGYAVMLAGLLYFILTIKNVDKKFQMS<br>NILSAVVMVSAFLLLYAQAQNWTSSFTFNEEVGRYFLDPSGDLFNNGYRYLNWLINVPML<br>LFQILFVVSLTTSKFSSVRNQFWFSGAMMIITGYIGQFYEVSNLTAFLVWGAISSAFFFH<br>ILWVMKKVINEGKEGISPAGQKILSNIWILFLISWTLYPGAYLMPYLTGVDGFLYSEDGV<br>MARQLVYTIADVSSKVIYGVLLGNLAITLSKNKIGTGFPFDPHYVEVLGERMHYVDVGPR<br>DGTPVLFLHGNPTSSYVWRNIIPHVAPTHRCIAPDLIGMGKSDKPDLGYFFDDHVRFMDA<br>FIEALGLEEVVLVIHDWGSALGFHWAKRNPERVKGIAFMEFIRPIPTWDEWPEFARETFQ<br>AFRTTDVGRKLIIDQNVFIEGTLPMGVVRPLTEVEMDHYREPFLNPVDREPLWRFPNELP<br>IAGEPANIVALVEEYMDWLHQSPVPKLLFWGTPGVLIPPAEAARLAKSLPNCKAVDIGPG<br>LNLLQEDNPDLIGSEIARWLSTLEISGEPTTKSRITSEGEYIPLDQIDINVFCYENEV |
| GvR (-) | MLMTVFSSAPELALLGSTFAQVDPSNLSVSDSLTYGQFNLVYNAFSFAIAAMFASALFFF<br>SAQALVGQRYRLALLVSAIVVSIAGYHYFRIFNSWDAAYVLENGVYSLTSEKFNDAYRYV<br>NWLLTVPLLLVETVAVLTLPAKEARPLLIKLTVASVLMIATGYPGEISDDITTRIIWGTV<br>STIPFAYILYVLWVELSRSLVRQPAAVQTLVRNMRWLLLLSWGVYPIAYLLPMLGVSGTS<br>AAVGVQVGYTIADVLAKPVFGLLVFAIALVKTKADIGTGFPFDPHYVEVLGERMHYVDVG<br>PRDGTPVLFLHGNPTSSYVWRNIIPHVAPTHRCIAPDLIGMGKSDKPDLGYFFDDHVRFM<br>DAFIEALGLEEVVLVIHDWGSALGFHWAKRNPERVKGIAFMEFIRPIPTWDEWPEFARET |

| | |
|---|---|
| | FQAFRTTDVGRKLIIDQNVFIEGTLPMGVVRPLTEVEMDHYREPFLNPVDREPLWRFPNE<br>LPIAGEPANIVALVEEYMDWLHQSPVPKLLFWGTPGVLIPPAEAARLAKSLPNCKAVDIG<br>PGLNLLQEDNPDLIGSEIARWLSTLEISGEPTTKSRITSEGEYIPLDQIDINVFCYENEV |
| SrR (-) | MLQELPTLTPGQYSLVFNMFSFTVATMTASFVFFVLARNNVAPKYRISMMVSALVVFIAG<br>YHYFRITSSWEAAYALQNGMYQPTGELFNDAYRYVNWLLTVPLLTVELVLVMGLPKNERG<br>PLAAKLGFLAALMIVLGYPGEVSENAALFGTRGLWGFLSTIPFVWILYILFTQLGDTIQR<br>QSSRVSTLLGNARLLLLATWGFYPIAYMIPMAFPEAFPSNTPGTIVALQVGYTIADVLAK<br>AGYGVLIYNIAKAKSEEEIGTGFPFDPHYVEVLGERMHYVDVGPRDGTPVLFLHGNPTSS<br>YVWRNIIPHVAPTHRCIAPDLIGMGKSDKPDLGYFFDDHVRFMDAFIEALGLEEVVLVIH<br>DWGSALGFHWAKRNPERVKGIAFMEFIRPIPTWDEWPEFARETFQAFRTTDVGRKLIIDQ<br>NVFIEGTLPMGVVRPLTEVEMDHYREPFLNPVDREPLWRFPNELPIAGEPANIVALVEEY<br>MDWLHQSPVPKLLFWGTPGVLIPPAEAARLAKSLPNCKAVDIGPGLNLLQEDNPDLIGSE<br>IARWLSTLEISGEPTTKSRITSEGEYIPLDQIDINVFCYENEV |
| AbR (-) | MAKPTVKEIKSLQNFNRIAGVFHLLQMLAVLALANDFALPMTGTYLNGPPGTTFSAPVVI<br>LETPVGLAVALFLGLSALFHFIVSSGNFFKRYSASLMKNQNIFRWVQYSLSSSVMIVLIA<br>QICGIADIVALLAIFGVNASMILFGWLQEKYTQPKDGDLLPFWFGCIAGIVPWIGLLIYV<br>IAPGSTSDVAVPGFVYGIIISLFLFFNSFALVQYLQYKGKGKWSNYLRGERAYIVLSLVA<br>KSALAWQIFSGTLIPAIGTGFPFDPHYVEVLGERMHYVDVGPRDGTPVLFLHGNPTSSYV<br>WRNIIPHVAPTHRCIAPDLIGMGKSDKPDLGYFFDDHVRFMDAFIEALGLEEVVLVIHDW<br>GSALGFHWAKRNPERVKGIAFMEFIRPIPTWDEWPEFARETFQAFRTTDVGRKLIIDQNV<br>FIEGTLPMGVVRPLTEVEMDHYREPFLNPVDREPLWRFPNELPIAGEPANIVALVEEYMD<br>WLHQSPVPKLLFWGTPGVLIPPAEAARLAKSLPNCKAVDIGPGLNLLQEDNPDLIGSEIA<br>RWLSTLEISGEPTTKSRITSEGEYIPLDQIDINVFCYENEV |
| HdR (-) | MSSHSATTSSGRRESARDSRLRLWNSVMAVLHFLQGAAMVLLADTVLWPITRTRYGFDPG<br>SQSIFPETVAFVDANLPLLVAGFLFISALAHTAIATVWYDKYVRYLDRGMNPYRWYEYSV<br>SASLMIVVIGMLSGVWDLGTLVALFGLVAVMNLSGLLMEQRNELTEQTDWTPYWVGVIAG<br>IVPWITIGVAFVGSVTASAGEFPEFVIYIYVSIFVFFNLFALNMALQYLEVSRWKNYLFG<br>EKMYIVLSLVAKSALAWQVYFGTLNSPIIGTGFPFDPHYVEVLGERMHYVDVGPRDGTPV<br>LFLHGNPTSSYVWRNIIPHVAPTHRCIAPDLIGMGKSDKPDLGYFFDDHVRFMDAFIEAL<br>GLEEVVLVIHDWGSALGFHWAKRNPERVKGIAFMEFIRPIPTWDEWPEFARETFQAFRTT<br>DVGRKLIIDQNVFIEGTLPMGVVRPLTEVEMDHYREPFLNPVDREPLWRFPNELPIAGEP<br>ANIVALVEEYMDWLHQSPVPKLLFWGTPGVLIPPAEAARLAKSLPNCKAVDIGPGLNLLQ<br>EDNPDLIGSEIARWLSTLEISGEPTTKSRITSEGEYIPLDQIDINVFCYENEV |
| TsR (-) | MTDYEDHTIRRFRIFNAIMGGIHLLQVFLVLYLSNNFSLPVTISKPVYNEFTNSISPVSE<br>TLFSIRVGPLVALFLFISAIAHILIATVLYYRYVENLKSGMNPYRWFEYSISASVMIVII<br>AMLTTIYDLGTLLALFTLTAVMNLMGLMMELHNQTTQNTDWTSYIIGCIAGLVPWIVIFI<br>PLIAAESVPDFVIYIFVSIAIFFNCFAINMYLQYKKIGKWKDYLHGERVYIILSLVAKSA<br>LAWQVFAGTLRPMIGTGFPFDPHYVEVLGERMHYVDVGPRDGTPVLFLHGNPTSSYVWRN<br>IIPHVAPTHRCIAPDLIGMGKSDKPDLGYFFDDHVRFMDAFIEALGLEEVVLVIHDWGSA<br>LGFHWAKRNPERVKGIAFMEFIRPIPTWDEWPEFARETFQAFRTTDVGRKLIIDQNVFIE<br>GTLPMGVVRPLTEVEMDHYREPFLNPVDREPLWRFPNELPIAGEPANIVALVEEYMDWLH<br>QSPVPKLLFWGTPGVLIPPAEAARLAKSLPNCKAVDIGPGLNLLQEDNPDLIGSEIARWL<br>STLEISGEPTTKSRITSEGEYIPLDQIDINVFCYENEV |
| EhR (-) | MDTIVLTINWFNFIAGLFHVALAIVCALLGDVSQTFKMYMPFTTFRNGTQENEVTIKYSG<br>YFPMTVIFVVYFSVTALFHMGNAFLWNDTYHRFLSNRKNPIRWTEYSITAPLMTAILAFI<br>AGSRNWLFVIAASTLTFASISVGFVLDEIRKYSFFVLGMIPFLVEFSLIILSLYLTTSCY<br>PSYLPITIFVEFGLWILFPFVSLLELSGINYKIGELVFIVLSFVSKSVLAIIILNSENVLK<br>NGVTGICIGTGFPFDPHYVEVLGERMHYVDVGPRDGTPVLFLHGNPTSSYVWRNIIPHVA<br>PTHRCIAPDLIGMGKSDKPDLGYFFDDHVRFMDAFIEALGLEEVVLVIHDWGSALGFHWA<br>KRNPERVKGIAFMEFIRPIPTWDEWPEFARETFQAFRTTDVGRKLIIDQNVFIEGTLPMG<br>VVRPLTEVEMDHYREPFLNPVDREPLWRFPNELPIAGEPANIVALVEEYMDWLHQSPVPK<br>LLFWGTPGVLIPPAEAARLAKSLPNCKAVDIGPGLNLLQEDNPDLIGSEIARWLSTLEIS<br>GEPTTKSRITSEGEYIPLDQIDINVFCYENEV |
| HaR (-) | MTKALEMTILDDGSESPISFKYLRKFNLAMGILHLVQGVAMLVLGFFWDFSRPLYSITLD<br>YSGGPPVAALQPEFSFTAVGPTVAAFLLFSALAHLLIAGPLNKFYVKNLKKKMNPIRWFE<br>YAFSSSIMVFFIAILFGVWDLWVLIGLFFLNMLMNLFGHMMELHNQTTKKTNWTAYIYGW<br>IAGIIPWVIISVFFARIAINSTGMPWFVPVIYAFELILFMSFAFNMLLQYKKVGKWKDYL<br>YGERMYQILSLVAKTLLAWLVFAGVFQPAIGTGFPFDPHYVEVLGERMHYVDVGPRDGTP<br>VLFLHGNPTSSYVWRNIIPHVAPTHRCIAPDLIGMGKSDKPDLGYFFDDHVRFMDAFIEA<br>LGLEEVVLVIHDWGSALGFHWAKRNPERVKGIAFMEFIRPIPTWDEWPEFARETFQAFRT<br>TDVGRKLIIDQNVFIEGTLPMGVVRPLTEVEMDHYREPFLNPVDREPLWRFPNELPIAGE<br>PANIVALVEEYMDWLHQSPVPKLLFWGTPGVLIPPAEAARLAKSLPNCKAVDIGPGLNLL<br>QEDNPDLIGSEIARWLSTLEISGEPTTKSRITSEGEYIPLDQIDINVFCYENEV |

| | |
|---|---|
| CR1 (+) | MPEPGSEAIWLWLGTAGMFLGMLYFIARGWGETDSRRQKFYIATILITAIAFVNYLAMAL<br>GFGLTIVEFAGEEHPIYWARYSDWLFTTPLLLYNLGLLAGADRNTITSLVSLDVLMIGTG<br>LVATLSPGSGVLSAGAERLVWWGISTAFLLVLLYFLFSSLSGRVADLPSDTRSTFKTLRN<br>LVTVVWLVYPVWWLIGTEGIGLVGIGIVTAGFMVIDLTAKVGFGIILLRSHGVLDGAAIG<br>TGFPFDPHYVEVLGERMHYVDVGPRDGTPVLFLHGNPTSSYVWRNIIPHVAPTHRCIAPD<br>LIGMGKSDKPDLGYFFDDHVRFMDAFIEALGLEEVVLVIHDWGSALGFHWAKRNPERVKG<br>IAFMEFIRPIPTWDEWPEFARETFQAFRTTDVGRKLIIDQNVFIEGTLPMGVVRPLTEVE<br>MDHYREPFLNPVDREPLWRFPNELPIAGEPANIVALVEEYMDWLHQSPVPKLLFWGTPGV<br>LIPPAEAARLAKSLPNCKAVDIGPGLNLLQEDNPDLIGSEIARWLSTLEISGEPTTKSRI<br>TSEGEYIPLDQIDINVFCYENEV |
| LmR (+) | MIVDQFEEVLMKTSQLFPLPTATQSAQPTHVAPVPTVLPDTPIYETVGDSGSKTLWVVFV<br>LMLIASAAFTALSWKIPVNRRLYHVITTIITLTAALSYFAMATGHGVALNKIVIRTQHDH<br>VPDTYETVYRQVYYARYIDWAITTPLLLLNLGLLAGMSGAHIFMAIVADLIMVLTGLFAA<br>FGSEGTPQKWGWYTIACIAYIFVVWHLVLNGGANARVKGEKLRSFFVAIGAYTLILWTAY<br>PIVWGLADGARKIGVDGVIIAYAVLDVLAKGVFGAWLLVTHANLRESDIGTGFPFDPHYV<br>EVLGERMHYVDVGPRDGTPVLFLHGNPTSSYVWRNIIPHVAPTHRCIAPDLIGMGKSDKP<br>DLGYFFDDHVRFMDAFIEALGLEEVVLVIHDWGSALGFHWAKRNPERVKGIAFMEFIRPI<br>PTWDEWPEFARETFQAFRTTDVGRKLIIDQNVFIEGTLPMGVVRPLTEVEMDHYREPFLN<br>PVDREPLWRFPNELPIAGEPANIVALVEEYMDWLHQSPVPKLLFWGTPGVLIPPAEAARL<br>AKSLPNCKAVDIGPGLNLLQEDNPDLIGSEIARWLSTLEISGEPTTKSRITSEGEYIPLD<br>QIDINVFCYENEV |
| PaR (+) | MIHHDQVAEMLYGYAGAAAAKPTHTDSPGPIPTVIPTPPQFQEIGETGHRTLWVVFALMV<br>LSSGFFAFMSWNVPISKRLYHVITTLITITASLSYFAMASGHVTSFSCTPAKDHHKHVPD<br>VGYTECRQVFWGRYVDWAITTPLLLLNLSLLAGIDGAHTLMAVIADVIMVLSGLFASQGE<br>TATQRWGWYAIGCVSYLFVIWHVALHGARTVTAKGRGVTRLFSSLALFTFVLWTAYPIVW<br>GIADGAHRTTVDTVILIYAVLDILAKPVFGLWLLFSHRSLAETNIGTGFPFDPHYVEVLG<br>ERMHYVDVGPRDGTPVLFLHGNPTSSYVWRNIIPHVAPTHRCIAPDLIGMGKSDKPDLGY<br>FFDDHVRFMDAFIEALGLEEVVLVIHDWGSALGFHWAKRNPERVKGIAFMEFIRPIPTWD<br>EWPEFARETFQAFRTTDVGRKLIIDQNVFIEGTLPMGVVRPLTEVEMDHYREPFLNPVDR<br>EPLWRFPNELPIAGEPANIVALVEEYMDWLHQSPVPKLLFWGTPGVLIPPAEAARLAKSL<br>PNCKAVDIGPGLNLLQEDNPDLIGSEIARWLSTLEISGEPTTKSRITSEGEYIPLDQIDI<br>NVFCYENEV |
| NcR (+) | MIHPEQVADMLRPTTSTTSSHVPGPVPTVVPTPTEYQTLGETGHRTLWVTFALMVLSSGI<br>FALLSWNVPTSKRLFHVITTLITVVASLSYFAMATGHATTFNCDTAWDHHKHVPDTSHQV<br>CRQVFWGRYVDWALTTPLLLLQLCLLAGVDGAHTLMAIVADVIMVLCGLFAALGEGGNTA<br>QKWGWYTIGCFSYLFVIWHVALHGSRTVTAKGRGVSRLFTGLAVFALLLWTAYPIIWGIA<br>GGARRTNVDTVILIYTVLDLLAKPVFGFWLLLSHRAMPETNIGTGFPFDPHYVEVLGERM<br>HYVDVGPRDGTPVLFLHGNPTSSYVWRNIIPHVAPTHRCIAPDLIGMGKSDKPDLGYFFD<br>DHVRFMDAFIEALGLEEVVLVIHDWGSALGFHWAKRNPERVKGIAFMEFIRPIPTWDEWP<br>EFARETFQAFRTTDVGRKLIIDQNVFIEGTLPMGVVRPLTEVEMDHYREPFLNPVDREPL<br>WRFPNELPIAGEPANIVALVEEYMDWLHQSPVPKLLFWGTPGVLIPPAEAARLAKSLPNC<br>KAVDIGPGLNLLQEDNPDLIGSEIARWLSTLEISGEPTTKSRITSEGEYIPLDQIDINVF<br>CYENEV |
| Ace1 (+) | MSNPNPFQTTLGTDAQWVVFAVMALAAIVFSIAVQFRPLPLRLTYYVNIAICTIAATAYY<br>AMAVNGGDNKPTAGTGADERQVIYARYIDWVFTTPLLLLNLVLLTNMPATMIAWIMGADI<br>AMIAFGIIGAFTVGSYKWFYFVVGCIMLAVLAWGMINPIFKEELQKHKEYTGAYTTLLIY<br>LIVLWVIYPIVWGLGAGGHIIGVDVVIIAMGVLDLLAKPLYAIGVLITVEVVYGKIGTGF<br>PFDPHYVEVLGERMHYVDVGPRDGTPVLFLHGNPTSSYVWRNIIPHVAPTHRCIAPDLIG<br>MGKSDKPDLGYFFDDHVRFMDAFIEALGLEEVVLVIHDWGSALGFHWAKRNPERVKGIAF<br>MEFIRPIPTWDEWPEFARETFQAFRTTDVGRKLIIDQNVFIEGTLPMGVVRPLTEVEMDH<br>YREPFLNPVDREPLWRFPNELPIAGEPANIVALVEEYMDWLHQSPVPKLLFWGTPGVLIP<br>PAEAARLAKSLPNCKAVDIGPGLNLLQEDNPDLIGSEIARWLSTLEISGEPTTKSRITSE<br>GEYIPLDQIDINVFCYENEV |
| CvR (+) | MAVHQIGEGGLVMYWVTFGLMAFSALAFAVMTFTRPLNKRSHGYITLAIVTIAAIAYYAM<br>AASGGKALVSNPDGNLRDIYYARYIDWFFTTPLLLLNIILLTGIPIGVTLWIVLADVAMI<br>MLGLFGALSTNSYRWGYYGVSCAFFFVVLWGLFFPGAKGARARGGQVPGLYFGLAGYLAL<br>LWFGYPIVWGLAEGSDYISVTAVAASYAGLDIAAKVVFGWAVMLSHPLIARNQIGTGFPF<br>DPHYVEVLGERMHYVDVGPRDGTPVLFLHGNPTSSYVWRNIIPHVAPTHRCIAPDLIGMG<br>KSDKPDLGYFFDDHVRFMDAFIEALGLEEVVLVIHDWGSALGFHWAKRNPERVKGIAFME<br>FIRPIPTWDEWPEFARETFQAFRTTDVGRKLIIDQNVFIEGTLPMGVVRPLTEVEMDHYR<br>EPFLNPVDREPLWRFPNELPIAGEPANIVALVEEYMDWLHQSPVPKLLFWGTPGVLIPPA |

| | |
|---|---|
| | EAARLAKSLPNCKAVDIGPGLNLLQEDNPDLIGSEIARWLSTLEISGEPTTKSRITSEGE YIPLDQIDINVFCYENEV |
| KnR (+) | MVYSHADAAGWTLYWITYGIMAVTALIFFAMSLRRPIQQRSHHYTSFLIVAIASLAYYAM ASQGGNTRIRVYQGPADSYRQIFWARYVDWFFTTPLLLLNLVLLSNLSKLRIAAIMVADI FMILTGLFGAVEARSNKWGWFVFGCIFMLYIFYELLVNVRKGAYARGGQHGMLYSVLLVW LLILWVQYPVVWGLAEGSSTVSSDTVIAWYAALDICAKCVFGFILLLGIESIDRKRIGTG FPFDPHYVEVLGERMHYVDVGPRDGTPVLFLHGNPTSSYVWRNIIPHVAPTHRCIAPDLI GMGKSDKPDLGYFFDDHVRFMDAFIEALGLEEVVLVIHDWGSALGFHWAKRNPERVKGIA FMEFIRPIPTWDEWPEFARETFQAFRTTDVGRKLIIDQNVFIEGTLPMGVVRPLTEVEMD HYREPFLNPVDREPLWRFPNELPIAGEPANIVALVEEYMDWLHQSPVPKLLFWGTPGVLI PPAEAARLAKSLPNCKAVDIGPGLNLLQEDNPDLIGSEIARWLSTLEISGEPTTKSRITS EGEYIPLDQIDINVFCYENEV |
| TdR (+) | MSSLFEKRNTAVATNYRGPTNSIVINEAGSDWYWAVFSVMAASAIVFSVMAAMTPRGERV FHYLTIAIVSVASVAYFTMAADLGSVAIISEFANYSSLPTRQVFYARYIDWVITTPLLLT NLMLLAGLPWSTIIFTIVMDEVMVLTGLFGAITPSSYKWGYFTFGMVAYFFVAWVLIVEA RKNAHRLGSDVHRLYIGIAIWTATLWTLYPVAWGLSEGGNVTSSDGVAIFYGVLDLLAKP VFGLWILLGHKGIGMDRIGTGFPFDPHYVEVLGERMHYVDVGPRDGTPVLFLHGNPTSSY VWRNIIPHVAPTHRCIAPDLIGMGKSDKPDLGYFFDDHVRFMDAFIEALGLEEVVLVIHD WGSALGFHWAKRNPERVKGIAFMEFIRPIPTWDEWPEFARETFQAFRTTDVGRKLIIDQN VFIEGTLPMGVVRPLTEVEMDHYREPFLNPVDREPLWRFPNELPIAGEPANIVALVEEYM DWLHQSPVPKLLFWGTPGVLIPPAEAARLAKSLPNCKAVDIGPGLNLLQEDNPDLIGSEI ARWLSTLEISGEPTTKSRITSEGEYIPLDQIDINVFCYENEV |
| EsR (+) | MESNLVRRNGALTVNTMTQNNQSVAIHQTVRGSDWYYTVCAVMGTTSLAILALSRMKPRT DRVFFYLTSGLCMVACIAYFAMGSNLGWTPIDVEWLRNDSVVRGVNRQVFYARYIDWVIT TPMLLMNLLLTAGMPWPTILWIILLDEIMIVTGLIGALVKSRYKWGFYVFGCMAMFYIMW ELAFPARKHAKVLGKDIHRSFVLCGVLTLVVWLCYPICWGLSEGGNVISPDSVSVFYGVL DVLAKPSFSIALIATHWNIDPGRIGTGFPFDPHYVEVLGERMHYVDVGPRDGTPVLFLHG NPTSSYVWRNIIPHVAPTHRCIAPDLIGMGKSDKPDLGYFFDDHVRFMDAFIEALGLEEV VLVIHDWGSALGFHWAKRNPERVKGIAFMEFIRPIPTWDEWPEFARETFQAFRTTDVGRK LIIDQNVFIEGTLPMGVVRPLTEVEMDHYREPFLNPVDREPLWRFPNELPIAGEPANIVA LVEEYMDWLHQSPVPKLLFWGTPGVLIPPAEAARLAKSLPNCKAVDIGPGLNLLQEDNPD LIGSEIARWLSTLEISGEPTTKSRITSEGEYIPLDQIDINVFCYENEV |
| MlR (+) | MGNSAIEVNGDTVDSYTADVKITTHGSDAYWAITAVMAFTTILFIAHSFTKPRTDRIFHY ITISITLVASIAYFTMASNLGWASIFIEFQRDDPLVSGTTREIFYVRYIDWVITTPLLLL NILLTAGLPWPTILFTILLDEVMIITGLVGALVKSSYKWGFFTFGCVAFFGVAWSVAWTG RKHANALGSDIGRVYLMTSVWTLFLWLLYPIAWGVSEGGNVISPDSVAAFYGTLDVLAKP IFGIILLWGHRNIPASRIGTGFPFDPHYVEVLGERMHYVDVGPRDGTPVLFLHGNPTSSY VWRNIIPHVAPTHRCIAPDLIGMGKSDKPDLGYFFDDHVRFMDAFIEALGLEEVVLVIHD WGSALGFHWAKRNPERVKGIAFMEFIRPIPTWDEWPEFARETFQAFRTTDVGRKLIIDQN VFIEGTLPMGVVRPLTEVEMDHYREPFLNPVDREPLWRFPNELPIAGEPANIVALVEEYM DWLHQSPVPKLLFWGTPGVLIPPAEAARLAKSLPNCKAVDIGPGLNLLQEDNPDLIGSEI ARWLSTLEISGEPTTKSRITSEGEYIPLDQIDINVFCYENEV |
| ScR (+) | MDSLLLQRRNDAVATNPPNATFALTENGSSWLWAVFCVMALSCIIIAALSLFKPMGYRIF YLLNVAILATASVSYFSLASDLGLTPVTVEFRGPGTRQIAYVRYIDWFVTTPLLLTQLLL TAGLPTNIIISTIFADLVMIITGLAGALVVSRYKWGYYTMGCVAMLWVFWNVFTGIKVSG NIGPDVRKSYTLLAVWLMIIWLNYPICWGLAEGGNRITVVGVMVYYGVLDLLAKPVFAAI SLAVHSKIELSRIGTGFPFDPHYVEVLGERMHYVDVGPRDGTPVLFLHGNPTSSYVWRNI IPHVAPTHRCIAPDLIGMGKSDKPDLGYFFDDHVRFMDAFIEALGLEEVVLVIHDWGSAL GFHWAKRNPERVKGIAFMEFIRPIPTWDEWPEFARETFQAFRTTDVGRKLIIDQNVFIEG TLPMGVVRPLTEVEMDHYREPFLNPVDREPLWRFPNELPIAGEPANIVALVEEYMDWLHQ SPVPKLLFWGTPGVLIPPAEAARLAKSLPNCKAVDIGPGLNLLQEDNPDLIGSEIARWLS TLEISGEPTTKSRITSEGEYIPLDQIDINVFCYENEV |
| RgR (+) | MDAILSKRNEVLSLNPLVANIDITTAASDWLWAVFAVMGLSAIILLVLGHATRPIGERAF HELAAAALCFTASIAYYSMASDLGATPIEVEFIRGGTLGQNWVDIGVLRPTRSIWYARYID WTITTPLLLLQLALTTALPLSQIFGLVFFDIVMIITGLLGALTASRYKWGFFVFGCVAMF WIFWVLFFPARKSASHLGTDYHRAYTSSAIVLCTLWTVYPIIWGVCDGGNVITPTSVMVA YGVLDLLAKPVFSFWHVFQLSRLDYARIGTGFPFDPHYVEVLGERMHYVDVGPRDGTPVL FLHGNPTSSYVWRNIIPHVAPTHRCIAPDLIGMGKSDKPDLGYFFDDHVRFMDAFIEALG LEEVVLVIHDWGSALGFHWAKRNPERVKGIAFMEFIRPIPTWDEWPEFARETFQAFRTTD VGRKLIIDQNVFIEGTLPMGVVRPLTEVEMDHYREPFLNPVDREPLWRFPNELPIAGEPA NIVALVEEYMDWLHQSPVPKLLFWGTPGVLIPPAEAARLAKSLPNCKAVDIGPGLNLLQE DNPDLIGSEIARWLSTLEISGEPTTKSRITSEGEYIPLDQIDINVFCYENEV |
| RtR (+) | MPFYIKNADIDITTHGSDWLWAVFSVMLLSAIGILVWGHVARPLGERAFHELAAALCFTA |

| | |
|---|---|
| | SIAYFAMASDLGDVPIVVEFIRGGSLGQNWVQVGVENPTRAIWYARYIDWTITTPMLLLQ<br>LLLCTGLPLSQVFSVIFADLLMIETGLIGALVASRYKWGFYAFGCAAQLYIWWMLLVPGR<br>RSAQHIGSDFAKSYTMSNIFLTTVWLVYPVIWGVADGGNVITPDSVMIAYGVLDLLAKPV<br>FSVIHLMSLSKLDYARIGTGFPFDPHYVEVLGERMHYVDVGPRDGTPVLFLHGNPTSSYV<br>WRNIIPHVAPTHRCIAPDLIGMGKSDKPDLGYFFDDHVRFMDAFIEALGLEEVVLVIHDW<br>GSALGFHWAKRNPERVKGIAFMEFIRPIPTWDEWPEFARETFQAFRTTDVGRKLIIDQNV<br>FIEGTLPMGVVRPLTEVEMDHYREPFLNPVDREPLWRFPNELPIAGEPANIVALVEEYMD<br>WLHQSPVPKLLFWGTPGVLIPPAEAARLAKSLPNCKAVDIGPGLNLLQEDNPDLIGSEIA<br>RWLSTLEISGEPTTKSRITSEGEYIPLDQIDINVFCYENEV |
| KeR1 (+) | MKFLLLLLADPTKLDPSDYVGFTFFVGAMAMMAASAFFFLSLNQFNKKWRTSVLVSGLIT<br>FIAAVHYWYMRDYWFAIQESPTFFRYVDWVLTVPLMCVQFYLILKVAGAKPALMWKLILF<br>SVIMLVTGYFGEAVFQDQAALWGAISGAAYFYIVYEIWLGSAKKLAVAAGGDILKAHKIL<br>CWFVLVGWAIYPLGYMLGTDGWYTSILGKGSVDVAYNIADAINKIGFGLVIYALAVKKNE<br>VDIGTGFPFDPHYVEVLGERMHYVDVGPRDGTPVLFLHGNPTSSYVWRNIIPHVAPTHRC<br>IAPDLIGMGKSDKPDLGYFFDDHVRFMDAFIEALGLEEVVLVIHDWGSALGFHWAKRNPE<br>RVKGIAFMEFIRPIPTWDEWPEFARETFQAFRTTDVGRKLIIDQNVFIEGTLPMGVVRPL<br>TEVEMDHYREPFLNPVDREPLWRFPNELPIAGEPANIVALVEEYMDWLHQSPVPKLLFWG<br>TPGVLIPPAEAARLAKSLPNCKAVDIGPGLNLLQEDNPDLIGSEIARWLSTLEISGEPTT<br>KSRITSEGEYIPLDQIDINVFCYENEV |
| KeR2 (+) | MTQELGNANFENFIGATEGFSEIAYQFTSHILTLGYAVMLAGLLYFILTIKNVDKKFQMS<br>NILSAVVMVSAFLLLYAQAQNWTSSFTFNEEVGRYFLDPSGDLFNNGYRYLNWLIDVPML<br>LFQILFVVSLTTSKFSSVRNQFWFSGAMMIITGYIGQFYEVSNLTAFLVWGAISSAFFFH<br>ILWVMKKVINEGKEGISPAGQKILSNIWILFLISWTLYPGAYLMPYLTGVDGFLYSEDGV<br>MARQLVYTIANVSSKVIYGVLLGNLAITLSKNKIGTGFPFDPHYVEVLGERMHYVDVGPR<br>DGTPVLFLHGNPTSSYVWRNIIPHVAPTHRCIAPDLIGMGKSDKPDLGYFFDDHVRFMDA<br>FIEALGLEEVVLVIHDWGSALGFHWAKRNPERVKGIAFMEFIRPIPTWDEWPEFARETFQ<br>AFRTTDVGRKLIIDQNVFIEGTLPMGVVRPLTEVEMDHYREPFLNPVDREPLWRFPNELP<br>IAGEPANIVALVEEYMDWLHQSPVPKLLFWGTPGVLIPPAEAARLAKSLPNCKAVDIGPG<br>LNLLQEDNPDLIGSEIARWLSTLEISGEPTTKSRITSEGEYIPLDQIDINVFCYENEV |
| GvR (+) | MLMTVFSSAPELALLGSTFAQVDPSNLSVSDSLTYGQFNLVYNAFSFAIAAMFASALFFF<br>SAQALVGQRYRLALLVSAIVVSIAGYHYFRIFNSWDAAYVLENGVYSLTSEKFNDAYRYV<br>DWLLTVPLLLVQTVAVLTLPAKEARPLLIKLTVASVLMIATGYPGEISDDITTRIIWGTV<br>STIPFAYILYVLWVELSRSLVRQPAAVQTLVRNMRWLLLLSWGVYPIAYLLPMLGVSGTS<br>AAVGVQVGYTIADVLAKPVFGLLVFAIALVKTKADIGTGFPFDPHYVEVLGERMHYVDVG<br>PRDGTPVLFLHGNPTSSYVWRNIIPHVAPTHRCIAPDLIGMGKSDKPDLGYFFDDHVRFM<br>DAFIEALGLEEVVLVIHDWGSALGFHWAKRNPERVKGIAFMEFIRPIPTWDEWPEFARET<br>FQAFRTTDVGRKLIIDQNVFIEGTLPMGVVRPLTEVEMDHYREPFLNPVDREPLWRFPNE<br>LPIAGEPANIVALVEEYMDWLHQSPVPKLLFWGTPGVLIPPAEAARLAKSLPNCKAVDIG<br>PGLNLLQEDNPDLIGSEIARWLSTLEISGEPTTKSRITSEGEYIPLDQIDINVFCYENEV |
| SrR (+) | MLQELPTLTPGQYSLVFNMFSFTVATMTASFVFFVLARNNVAPKYRISMMVSALVVFIAG<br>YHYFRITSSWEAAYALQNGMYQPTGELFNDAYRYVDWLLTVPLLTVQLVLVMGLPKNERG<br>PLAAKLGFLAALMIVLGYPGEVSENAALFGTRGLWGFLSTIPFVWILYILFTQLGDTIQR<br>QSSRVSTLLGNARLLLLATWGFYPIAYMIPMAFPEAFPSNTPGTIVALQVGYTIADVLAK<br>AGYGVLIYNIAKAKSEEEIGTGFPFDPHYVEVLGERMHYVDVGPRDGTPVLFLHGNPTSS<br>YVWRNIIPHVAPTHRCIAPDLIGMGKSDKPDLGYFFDDHVRFMDAFIEALGLEEVVLVIH<br>DWGSALGFHWAKRNPERVKGIAFMEFIRPIPTWDEWPEFARETFQAFRTTDVGRKLIIDQ<br>NVFIEGTLPMGVVRPLTEVEMDHYREPFLNPVDREPLWRFPNELPIAGEPANIVALVEEY<br>MDWLHQSPVPKLLFWGTPGVLIPPAEAARLAKSLPNCKAVDIGPGLNLLQEDNPDLIGSE<br>IARWLSTLEISGEPTTKSRITSEGEYIPLDQIDINVFCYENEV |
| PsuCCR (+) | MTMLEHLEGTMDGWYAENDLGQGAIIAHWVTFFFHMITTFYLGYVSFHSKGPGGKQPYFA<br>GYHEENNIGIFVNLFAAISYFGKVVSDTHGHNYQNVGPFIIGLGNYRYADYMLTCPLLVM<br>NLLFQLRAPYKITCAMLIFAVLMIGAVTNFYPGDDMKGPAVAWFCFGCFWYLIAYIFMAH<br>IVSKQYGRLDYLAHGTKAEGALFSLKLAIITFFAIWVAFPLVWLLSVGTGVLSNEAAEIC<br>HCICDVVAKSVYGFALANFREQYDRELIGTGFPFDPHYVEVLGERMHYVDVGPRDGTPVL<br>FLHGNPTSSYVWRNIIPHVAPTHRCIAPDLIGMGKSDKPDLGYFFDDHVRFMDAFIEALG<br>LEEVVLVIHDWGSALGFHWAKRNPERVKGIAFMEFIRPIPTWDEWPEFARETFQAFRTTD<br>VGRKLIIDQNVFIEGTLPMGVVRPLTEVEMDHYREPFLNPVDREPLWRFPNELPIAGEPA<br>NIVALVEEYMDWLHQSPVPKLLFWGTPGVLIPPAEAARLAKSLPNCKAVDIGPGLNLLQE<br>DNPDLIGSEIARWLSTLEISGEPTTKSRITSEGEYIPLDQIDINVFCYENEV |
| GtCCR1 (+) | MVESSAVIAANWISFLVIAGSFVVLCFISLRYKGPGGNENYYNGFREQNMLTVIINLWCA<br>LAYFAKVLQSHSDDDGFVPLTKIPYLDYATTCPLLTLNLMWCLDAPYKITSAVLVFTVMI<br>TGVACSLAVAPYSFYWFAMGMVLFIFTYVLMLSIVRERLEFITQCAHDSNAKRSIKHLKA<br>AVIIYFGIWPIFAILWLLSYRAANVISNDTNHILHCILDVIAKSCFGFVLLHFKMYFDKK |

| | |
|---|---|
| | LIGTGFPFDPHYVEVLGERMHYVDVGPRDGTPVLFLHGNPTSSYVWRNIIPHVAPTHRCI<br>APDLIGMGKSDKPDLGYFFDDHVRFMDAFIEALGLEEVVLVIHDWGSALGFHWAKRNPER<br>VKGIAFMEFIRPIPTWDEWPEFARETFQAFRTTDVGRKLIIDQNVFIEGTLPMGVVRPLT<br>EVEMDHYREPFLNPVDREPLWRFPNELPIAGEPANIVALVEEYMDWLHQSPVPKLLFWGT<br>PGVLIPPAEAARLAKSLPNCKAVDIGPGLNLLQEDNPDLIGSEIARWLSTLEISGEPTTK<br>SRITSEGEYIPLDQIDINVFCYENEV |
| GtCCR2 (+) | MVASSAVITANWISFLAISASFIILLVISLRYKGPGGTESFYNGFKEQNMLTVFINLWCA<br>LAYFAKVLQSHSNDNGFAPLTVIPYVDYCTTCPLLTLNLLWCLDAPYKISSAVLVFTCLV<br>IAVACSLAVAPFSYCWFAMGMVLFTFTYVFILSIVRQRLDFFTLCARDSNAKQSLKHLKT<br>AVFIYFGIWLLFPLLWLLSYRAANVISNDINHIFHCILDVIAKSVYGFALLYFKMYFDKK<br>LIGTGFPFDPHYVEVLGERMHYVDVGPRDGTPVLFLHGNPTSSYVWRNIIPHVAPTHRCI<br>APDLIGMGKSDKPDLGYFFDDHVRFMDAFIEALGLEEVVLVIHDWGSALGFHWAKRNPER<br>VKGIAFMEFIRPIPTWDEWPEFARETFQAFRTTDVGRKLIIDQNVFIEGTLPMGVVRPLT<br>EVEMDHYREPFLNPVDREPLWRFPNELPIAGEPANIVALVEEYMDWLHQSPVPKLLFWGT<br>PGVLIPPAEAARLAKSLPNCKAVDIGPGLNLLQEDNPDLIGSEIARWLSTLEISGEPTTK<br>SRITSEGEYIPLDQIDINVFCYENEV |
| GtCCR3 (+) | MVSALDQNGPQYLQNPIVIAADWIGFIALFGSSLAVAYKLVTFKGPDQDDVYFFGYREEK<br>MISVFVNLFAALAYWAKLASHANGDVGPAASVTTYKYLDYLFTCPLLTINLLWCLNLPYK<br>FTFGAIVAVCILCAFMASVIPPPARYMWFGMGITVFSAAWFNILKLVRMRLEQFVSKEAK<br>KVRQSLKVACMTYFFIWLGYPTLWVLGDAGVLDSVVSALLHTFLDVFSKSIYGFALLHFV<br>MRTDKREIGTGFPFDPHYVEVLGERMHYVDVGPRDGTPVLFLHGNPTSSYVWRNIIPHVA<br>PTHRCIAPDLIGMGKSDKPDLGYFFDDHVRFMDAFIEALGLEEVVLVIHDWGSALGFHWA<br>KRNPERVKGIAFMEFIRPIPTWDEWPEFARETFQAFRTTDVGRKLIIDQNVFIEGTLPMG<br>VVRPLTEVEMDHYREPFLNPVDREPLWRFPNELPIAGEPANIVALVEEYMDWLHQSPVPK<br>LLFWGTPGVLIPPAEAARLAKSLPNCKAVDIGPGLNLLQEDNPDLIGSEIARWLSTLEIS<br>GEPTTKSRITSEGEYIPLDQIDINVFCYENEV |
| GtCCR4 (+) | MTTSAPSLSDPNWQYGMGGWNNPRLPNFNLHDPTVIGVDWLGFLCLLGASLALMYKLMSF<br>KGPDGDQEFFVGYREEKCLSIYVNLIAAITYWGRICAHFNNDMGLSLSVNYFKYLDYIFT<br>CPILTLNLLWSLNLPYKITYSLFVGLTIACNAFEPPARYLWFMFGCFIFAFTWISIIRLV<br>YARFQQFLNEDAKKIRAPLKLSLTLYFSIWCGYPALWLLTEFGAISQLAAHVMTVIMDVA<br>AKSVYGFALLKFQLGVDKRDIGTGFPFDPHYVEVLGERMHYVDVGPRDGTPVLFLHGNPT<br>SSYVWRNIIPHVAPTHRCIAPDLIGMGKSDKPDLGYFFDDHVRFMDAFIEALGLEEVVLV<br>IHDWGSALGFHWAKRNPERVKGIAFMEFIRPIPTWDEWPEFARETFQAFRTTDVGRKLII<br>DQNVFIEGTLPMGVVRPLTEVEMDHYREPFLNPVDREPLWRFPNELPIAGEPANIVALVE<br>EYMDWLHQSPVPKLLFWGTPGVLIPPAEAARLAKSLPNCKAVDIGPGLNLLQEDNPDLIG<br>SEIARWLSTLEISGEPTTKSRITSEGEYIPLDQIDINVFCYENEV |
| GtCCR5 (+) | MSTSSVAYLRTPVVQALDWVGFISLGGTAAVLAYRLMNFKPPNKDILYFFGYREKGMISL<br>YVNLFAAVAYYARITSHLSGDVGAATNIILYKYFDYLITCPLLTFNLLTTLNLPYKITYA<br>VYVQITIFTGFMSANTPPPATFLWFAFGMLLFSYTWFNIISLVQVRFIQYFAKKGNTTQS<br>RRVSVASKAGFRNKNVRNPLQTALSTYFCIWMVYPVLWLLLKTKVIDQVTEHCINVVMDV<br>LAKSMYGFALLRFQLLMDKANIGTGFPFDPHYVEVLGERMHYVDVGPRDGTPVLFLHGNP<br>TSSYVWRNIIPHVAPTHRCIAPDLIGMGKSDKPDLGYFFDDHVRFMDAFIEALGLEEVVL<br>VIHDWGSALGFHWAKRNPERVKGIAFMEFIRPIPTWDEWPEFARETFQAFRTTDVGRKLI<br>IDQNVFIEGTLPMGVVRPLTEVEMDHYREPFLNPVDREPLWRFPNELPIAGEPANIVALV<br>EEYMDWLHQSPVPKLLFWGTPGVLIPPAEAARLAKSLPNCKAVDIGPGLNLLQEDNPDLI<br>GSEIARWLSTLEISGEPTTKSRITSEGEYIPLDQIDINVFCYENEV |

**Table 3. Summary of voltage optical recording conditions**

| Illumination | Microscope | Light source | Light power |
|---|---|---|---|
| 1-Photon | 1P widefield (Nikon Ti2) | LEDs (Spectra X light engine, Lumencore) | 10-50 $mW/mm^2$ |
| 2-Photon | 2P SLAP microscope | 1030 nm laser (Yb:YAG, 1,030 nm, 190 fs, tunable repetition rate 1–10 MHz; BlueCut, Menlo Systems) | 40 mW out of objective |
| 1-Photon | 1P widefield | 530 nm LED light (Thorlabs) | 10-50 $mW/mm^2$ |
| 2-Photon | Laser circularly scanned the neurons at 1 KHz | Monaco 1035 nm laser, repetition rate set to 10 MHz | 25.7 mW out of objective |
| 2-Photon | Laser patterned into a stationary broadened focal spot | Monaco 1035 nm laser, repetition rate set to 10 MHz | 50 mW out of objective |
| 2-Photon | 2P AOD microscope | InSight X3, Spectra Physics mode-locked at 1045nm with a repetition rate of 80 MHz | 15 mW out of objective |
