## Supplementary Materials and Methods for "A novel rhodopsin-based voltage indicator for simultaneous two-photon optical recording with GCaMP in vivo"

**Reagent availability.** Plasmids have been deposited at Addgene (www.addgene.org) as follows: pCAG-2Photron-ST (plasmid #224359), pCAG-PaR-HaloTag-ST (plasmid # 224360), pAAV-syn-FLEX-2Photron-ST (plasmid # 224361).

**Molecular biology.** The genes for all opsins described were synthesized (Integrated DNA Technologies) with mammalian codon optimization. HaloTag was amplified from the Voltron plasmid (addgene plasmid# 119033). Rhodopsins and HaloTag were combined using overlap PCR. Cloning was done by restriction enzyme digest of plasmid backbones, PCR amplification of inserted genes, isothermal assembly, and followed by Sanger sequencing to verify DNA sequences. A soma localization tag (ST) was added using overlap PCR to make soma targeted versions of indicators. For expression in primary neuron cultures, sensors were cloned into a pcDNA3.1-CAG plasmid (Invitrogen). The amino acid sequences of all tested opsin-HaloTag fusions are given in Supplementary Fig. 5 and Table 1 and 2. For the preparation of viruses, plasmid DNA was purified by the Janelia Molecular Biology Facility and AAVs were prepared by the Janelia Viral Tools Team.

**Imaging of primary neuron cultures**. Primary rat hippocampal neurons were prepared and transfected as described previously[1,2]. Stock solutions of JF525 HaloTag ligand was prepared at 1 mM in DMSO. Cultured neurons were incubated at 37°C for 20-30 min at a final concentration ~500 nM before washing twice with imaging buffer containing 145 mM NaCl, 2.5 mM KCl, 10 mM glucose, 10 mM HEPES, pH 7.4, 2 mM $CaCl_2$ and 1 mM $MgCl_2$. Synaptic blockers (10 μM CNQX, 10 μM CPP, 10 μM GABAZINE, and 1 mM MCPG) were added prior to imaging. Wide-field 1-photon imaging was performed on an inverted Nikon Eclipse Ti2 microscope equipped with a SPECTRA X light engine (Lumencore) with a 40X objective (NA = 1.3, Nikon), and imaged onto a sCMOS camera (Hamamatsu ORCA-Flash 4.0). A custom filter set (510/25 nm (excitation, Lumencore), 545/40 nm (emission, Chroma), and a 525LP dichroic mirror (Chroma)) was used to image all opsin-HT fusions labeled with JF525 in culture.

**Field stimulation of primary neuron cultures.** A stimulus isolator (A385, World Precision Instruments) with platinum wires was used to deliver field stimuli (50V, 1 ms) to elicit action potentials in cultured neurons as described previously[1]. The stimulation was controlled using Wavesurfer and timing was synchronized with fluorescence acquisition using Wavesurfer and a National Instruments PCIe-6353 board. To measure fluorescence response to action potentials over time, neurons were labeled with JF525 HaloTag ligand and imaged with a 40x objective at 400 Hz. We applied field electrode stimulations to induce a train of 10 single action potentials at 66Hz, followed by a train of 10 action potentials at 10 Hz. Images were processed in ImageJ. Voltage-dependent fluorescence changes were measured for hand-segmented neurons, on 2-3 well replicates and multiple fields of view per well.

**Simultaneous electrophysiology and fluorescence imaging in primary neuron culture under both 1p and 2p illumination**

All imaging and electrophysiology measurements were performed in imaging buffer (145 mM NaCl, 2.5 mM KCl, 10 mM glucose, 10 mM HEPES, pH 7.4, 2 mM $CaCl_2$, 1 mM $MgCl_2$) adjusted to 310 mOsm with sucrose. For voltage clamp measurements, 500 nM TTX was added to the imaging buffer to block sodium channels. Synaptic blockers (10 μM CNQX, 10 μM CPP, 10 μM GABAZINE, and 1 mM MCPG) were added to block ionotropic glutamate, GABA, and metabotropic glutamate receptors[1].

Glass capillary with filament (Sutter Instruments) were pulled to a tip resistance of 4 – 6 MΩ. Internal solution for current clamp recordings contained the following: 130 mM potassium methanesulfonate, 10 mM HEPES, 5 mM NaCl, 1 mM MgCl2, 1 mM Mg-ATP, 0.4 mM Na-GTP, 14 mM Tris-phosphocreatine, adjusted to pH 7.3 with KOH, and adjusted to 300 mOsm with sucrose. Internal solution for voltage clamp recordings contained the following: 115 mM cesium methanesulfonate,10 mM HEPES, 5 mM NaF, 10 mM EGTA, 15 mM CsCl, 3.5 mM Mg-ATP, 3 mM QX-314, adjusted to pH 7.3 with CsOH, and adjusted to 300 mOsm with sucrose.

Pipettes were positioned with a MP-285 manipulator (Sutter Instruments). Whole cell voltage clamp and current clamp recordings were acquired using an Axon700B amplifier, filtered at 10 kHz with the internal Bessel filter, and digitized using a National Instruments PCIe-6353 acquisition board at 20 kHz. Data were acquired from cells with access resistance < 25 MΩ. WaveSurfer software was used to generate the various analog and digital waveforms to control the amplifier, camera, light source, and record voltage and current traces. For fluorescence voltage curves, cells were held at a potential of –70 mV at the start of each step and then 0.5 s voltage steps were applied to step the potential from –145 mV to +55 mV in 25 mV increments. For current-clamp recordings to generate action potentials, current was injected (60-200 pA for 0.5 s) and voltage was monitored.

To image cells, two setups were used. In Setup 1 (Fig. 2 and Supplementary Fig. 7, 8) a Bergamo microscope (Thorlabs) operated with ScanImage software (Vidrio) was configured for 1p widefield excitation with a 530 nm LED (Thorlabs), and also for 2p excitation at 1035 nm from a femtosecond laser (Monaco-40 Watt, Coherent) running at 10 MHz pulse rate. To match the 2p beam size to the ~10 um cell dimension, either the 2p optics were configured to give a spatially extended beam over a selected cell (Fig.2 and Supplementary Fig. 7), or the beam was passed through a galvo-galvo scanner to deliver a focused spot circularly scanning the cell at a rate of 1 kHz (Supplementary Fig 8). For all imaging, a 16x water-dipping objective (CF175, Nikon) was used, and the epifluorescence beam path included a 1030 dichroic to bring in the 2p beam, a 525 dichroic to bring in the 1p 530 LED light, a 607/70 emission bandpass filter. a 200 mm tube lens (Thorlabs), and a Neo CMOS camera (Andor) set at the focal plane of the tube lens. The camera was operated with Andor Solis software to collect fluorescent images at a frame rate of 500 or 1000 fps with a reduced sensor size of 128 x 128 pixels (52 x 52 $um^2$ in object space).

In Setup 2, for Supplementary Fig. 6, data were collected on an inverted Nikon Eclipse Ti2 microscope equipped with a SPECTRA X light engine (Lumencore) with a 40X objective (NA = 1.3, Nikon) and a 550/15 nm excitation filter. A dichroic mirror (89100bs; Chroma) and 605/52 nm emission filters (Chroma) were also used. Image acquisition rate is 3.2 kHz. Fluorescence intensity was quantified using ImageJ software and data analysis performed in MATLAB and Prism GraphPad.

**Simultaneous dual fluorescence optical recordings in cortical neurons in behaving mice.**

All protocols adhered to the guidelines of the French National Ethics Committee for Sciences and Health report on Ethical Principles for Animal Experimentation in agreement with the European Community Directive 86/609/EEC under agreement #29791.

**Viral vector construction and packaging**

The 2Photron-Ts-Halotag construct is described above and the Voltron2-Ts-Halotag construct has been published[3]. All GEVIs were expressed as CRE-dependent type 1 AAVs under the control of the human synapsin (hSyn) promoter except for JEDI2P in the cortex where EF1a was used. Injected titers were $3e10^{12}$ GC/ml in the visual cortex and $4e10^{12}$ GC/mL in the cerebellum. The CRE-dependent GECIs GCaMP6f (Addgene, 100833-AAV1) and GCaMP8f (Addgene, 162379-AAV1) under the control of hSyn promoter were used at final titers of $1e10^{12}$ and $8e10^{11}$ GC/ml, respectively. To drive the *cre* recombinase in the visual cortex, AAV2/1-hSyn-Cre (Addgene, AV-1-PV2676) was injected at a final concentration of 1.5-$2e10^{9}$ GC/ml. In the cerebellum, to target glycinergic granular layer neurons, we used heterozygous GlyT2-Cre mice (kind gift of H.U. Zeilhofer, University of Zurich).

**Animal handling, viral injections, and surgeries**

9 male GlyT2-Cre, 9 male and 1 female wild-type C57BL/6J mice were housed in standard conditions (12-hour light/dark cycles, light on at 7 a.m., with water and food *ad libitum*). A preoperative analgesic was used (buprenorphine, 0.1 mg/kg), and Ketamine-Xylazine were used as anesthetics. Viruses were combined in PBS, 300 n of which was injected at a flow rate of 75 nl/min into the visual cortex (V1 coordinates from bregma: anteroposterior -3/-3.5mm, mediolateral -2.5/-3 mm, and dorsoventral -0.3mm from brain surface, 10 mice) or into the lobule IV-V of the cerebellar cortex (coordinates from bregma: anteroposterior -6mm, mediolateral ±0.4 mm, and dorsoventral -0.32mm from brain surface, 9 mice), of adult (>P40) mice (body weight 24–30 g). A custom-designed aluminum head-plate was fixed on the skull with layers of dental cement (Metabond). A 5 mm diameter (visual cortex) or custom-designed laser cut (cerebellum) #1 coverslip was placed on top of the targeted area and secured with dental cement (tetric evoflow). Mice were allowed to recover for at least 15 days before recording sessions and housed 2-4 mice per cage. Behavioral habituation was adopted, involving progressive handling by the experimenter with gradual increases in head-fixed duration[4]. One or 2 days before experiments, JF552 was prepared and retro-orboritally injected as described previously[2].

Mice were handled before recording sessions to limit restraint-associated stress, and experiments were performed during the light cycle.

**ULoVE voltage optical recording and experimental design**

3-4 hour recording sessions were performed while mice behaved spontaneously on top of an unconstrained running wheel in the dark[4]. Recordings were performed using a custom-designed dual scanner of acousto-optic deflector (AOD) -based random-access multiphoton microscope (Karthala System) based on a previously described design[5]. Excitation was provided by a femtosecond laser (InSight X3, Spectra Physics) mode-locked at 920 and 1045nm with a repetition rate of 80 MHz for a green or red optogenetic reporter, respectively.

A water-immersion objective (CFO Apo25XC W1300, 1.1 NA, 2 mm working distance, Nikon) was used for excitation and epifluorescence light collection. Laser power was set to deliver 15 mW post-objective and pre-sample, then adjusted for mono-exponential loss through tissue with a length constant of 170 µm. The power was further doubled to account for the greater excitation volume using ULoVE compared with that used in standard 2P laser scanning microscopy. Optimized ULoVE excitation pattern (Lombardini et al., in prep) consists of a series of 9 points vertically aligned (spread linearly over a distance of 15µm), multiplexed twice with a 2µm interpoint horizontal spacing and scanned during 50µs horizontally for a distance of 8 µm and 3µm vertically in order to keep the cell membrane in that excited volume. This new strategy enables to limit the axial recomposition, thus improving cell specific voltage SNR. Two or three of those optimized multiplexed patterns (4 patterns in total: red + green) were placed on the cell membrane for GEVI excitation (see supplementary Fig. 10c) and positioned on the cytoplasm for GECI excitation. To avoid crosstalk between channels during dual excitation recordings, transmission at the AOM was set to 0 on the 920nm laser line for recording red signals and vice versa on the 1045nm laser line for recording green signals. As added controls, patterns were positioned outside of the targeted neuron. The efficiency of this is presented in supplementary fig.10d. Using four patterns, temporal resolutions of 3751 Hz (visual cortex) and 3751- 4761 Hz (3 to 4 patterns) (cerebellum) were obtained. Signals were passed through an IR blocking filter (TF1, Thorlabs), split into two channels using a 562 nm dichroic mirror (Semrock), and passed to two H12056P-40 photomultiplier tubes (Hamamatsu) used in photon counting mode. A 510/84 filtered green channel was used for JEDI-2P and a 607/70 filtered red channel was used for the rhodopsin-based GEVIs. Cells chosen for recording were manually selected based on brightness sufficient to yield significant signal-to-noise (see supplementary Fig. 11f). Recordings were stopped after 10 min and repeated across multiple cells and mice (details in the figure legends).

**Photon flux analysis**

In the dual GEVI recording experiments of 2Photron + JEDI2P we observed an inverse relationship between baseline signals, such that we found no one cell displaying both GEVIs at the “normal” bright level seen when either was expressed singly. Thus, to quantify the photon flux of a large number of cells and to identify cells that were coexpressing the two GEVIs, we acquired 2P stacks (4 mice) at a 1 µs pixel dwell time, 5.5 pixels per micron, and 2 µm step size in z. Cells hand-selected within the stacks were then positioned in a 100 ± 25 pixel-sized region and the z plane centered at the cell equator. Half of the highest photon counts inside this box, ± one plane, were summed and the flux was expressed in MHz. As mentioned, the cells showing the brightest baseline JEDI-2P expression had weak baseline 2Photron brightness, so for the recordings shown in Fig. 3c-g, we had to compromise GEVI photon flux at the expense of the detectability supplementary Fig. 11d-f.

**Signal analyses**

Traces were generated by summing the collected photons per ULoVE volume and photobleaching was corrected by bi-exponential fitting. After removing low-frequency drift using a zero-phase distortion filter (highpass: 0.5 Hz), the traces were converted to % ∆F/F taking the mean signal as Fo. Up and down states were then assigned using a double Gaussian fit on the histogram distribution and the down state was subtracted. To compare the subthreshold between 2Photron and JEDI-2P (Fig. 3 c & d), traces were lowpass filtered at 25 Hz, and a linear regression between them was performed to extract the slope. For the calcium trace, a similar approach was performed but the baseline used to convert the trace to %∆F/F was the mode of the calcium trace photon count. For display purpose in figures, optical recording traces were smoothed with a Gaussian kernel of 0.25 ms. To compare photon flux and related discriminability metrics between ASAP3, JEDI-2P and 2Photron (fig. 3f), we had to normalize the values to account for changes in applied power, pattern number per cell and per channel and pattern efficiency compared to the original single cell and non-multiplexed ASAP3 related data such as correcting factor is 3.12 for JEDI-2P and 2.0 for 2Photron.

**Spikes extraction and waveform analyses.**

Spikes were detected with a custom-designed algorithm. We combined three metrics that encompass different aspects of spike shapes and thresholds, all with a z-score above 2.5 in at least two of the metrics, and 2 as a minimal level for the third metric. The first metric is the highpass filtered trace (second order Butterworth filter with the lower limit set at 40 Hz). The second metric is a cumulative probability transform of the signal using the erf function for a duration of 1.6 ms. The third metric is the cumulative product of the $2^{nd}$ to the $5^{th}$ scale of the coif1 discrete wavelet transform followed by a global realignment that enables energy retrieval in various frequency bands in one peak. After detection, all individual events were visually inspected in this wavelet domain, and any questionable detections were ruled out.

To perform the ground truth analysis (supplementary Fig. 11), the JEDI-2P detected spikes were taken as reference. Spikes falling within 3 ms of a given reference spike qualified as a true positive.

Sensitivity is defined as the ratio between the number of detected spikes and the number of reference spikes. Specificity is the ratio between false positives and the number of reference spikes. Precision is the ratio between the number of detected spikes and the number of all detected spikes. The error rate corresponds to 1 minus the harmonic mean of sensitivity and precision[6].

To compute spike waveform-related metrics, a spike average waveform based on the onset was computed where the amplitude at the onset was set to 0% ΔF/F. Amplitude corresponds to the peak value of the spike waveform average. FWHM corresponds to the extent in time at half of its amplitude after a 10KHz interpolation of the spike average waveform. Tau of the depolarization and repolarization were extracted from mono and bi-exponential fits of the average spike waveform, respectively. For cerebellar cells, AHP corresponds to the trough magnitude in the average waveform, and dynamic range is the absolute sum of the AHP and spike magnitude. Spike SNR is the spike amplitude (in photons) divided by the shot noise, corresponding to the square root of the mean photon count of the trace. D' is computed as previously reported[5,7].

Bursts were extracted as the group of spikes that have an inter-spike interval (isi) below a limit defined for each neuron (mean ± std: 25.8 +/- 9.69 ms). To define this limit, the isi distribution, with a 1ms time bin and in log-log scale, was fitted by a Gaussian curve and the burst duration limit corresponds to the duration at which the ordinate of the fit is 0.2.

To extract the metric that we call 2Photron signal at spike onset (Fig. 4 h, j, k), we first obtain the baseline drift from MLspike[6], which provides a value of this drift at each spike onset.

To extract the depolarization with and without spikes (Fig. 4l), first, we average the traces of all bursts above 3 spikes. We use this template to set the parameters of the differential transform of the trace, which provides enhanced contrast of such average depolarizations. We found that performing the bidirectional differential of the signal during a 15 ms period of up-state and a baseline of 50 ms, separated by a lag of 30 ms, generated peaks when depolarizations occurred in both traces.

We then set a detection z-score threshold of two, together with a prominence of one z-score, and sorted the detected events according to the presence or absence of depolarizations also bearing spikes. Events with spikes and calcium transients preceding the detected events were discarded.

### Calcium transient-related analyses

All bursts selected for subsequent analyses were kept if their preceding isi was larger than 50 ms and the traces were back to baseline level before the onset of the next burst. Calcium transients were extracted using as an onset the first spike of the burst. The amplitude of a calcium transient corresponds to the maximal value following the last spike of the burst, from a trace smoothed with a Gaussian kernel of 5 ms. GCaMP8f cells that strictly displayed more than 3 burst sizes were kept for subsequent analyses (9/12 cells).

To correct for inter-cell variance, we normalized the calcium transient amplitude at the single-cell level, either using realistic or scaled normalization. Realistic normalization tends to minimize differences between cells by normalizing amplitudes using the dynamic range (lowest-highest; aka 1 to 99 percentiles of the distribution) seen for that cell.

$$realistic\ normalized\ amplitude(cell) = \left(\frac{amplitudes(cell) - 1th\ prctile(cell)}{99th\ prctile\ (cell)}\right)$$

The scaled normalization is done in two steps, first, we compute the mean of calcium transient amplitude per burst size. We then fit a quadratic function using the following equation:

$$fit(cell) = \alpha\ x^2$$

Then we minimize the residual amplitude and adjust the b parameter for the whole cell population (b = 4.108 and 2.333 respectively for the 2Photron and Zhang datasets).

Then cells are refitted to determine the scaling factor $\alpha$ :

$$fit(cell) = \alpha_{cell}(x^2 + b)$$

and then we use this scaling factor for the normalization such as:

$$scaled\ normalized\ amplitude(cell) = \left(\frac{amplitudes(cell)}{\alpha_{cell}}\right)$$

To study calcium amplitude distributions as a function of burst size, the cumulative normalized calcium transient distributions (0.05 bin size) were fit using sigmoid functions for each burst size. The derivative of those fits is used to obtain the probability of event amplitudes knowing the burst size.

To compute true and false probabilities, we first computed the probability of obtaining an event of any burst size knowing the amplitude using the following:

$$p(burst|amp) = \frac{p(amp|burst) * p(burst)}{p(amp)}$$

Where $p(amp|burst)$ is the probability of event amplitudes knowing the burst size, $p(burst)$ is the fraction of bursts occurring amongst all bursts included in the analyses, and $p(amp)$ is the fraction of normalized amplitudes regardless of the burst size (obtained from the derivative of the sigmoid fits).

To find a true positive fraction, we first ascertained the true burst size for a given amplitude range simply by taking the burst size that had the highest probability distribution. The true positive amplitude corresponds to the fraction of events for this tested amplitude for which the true burst size is equal to the tested burst size. If the tested burst size is equal to the true burst size ± 1 spike/burst it falls in the false positive 1 spike error category and similarly for 2, 3, and up to ± 4 spikes/burst maximal error.

To provide a single value for true and errors over the range of tested amplitudes, we computed the mean values of those true and false positive fractions weighted by the distribution probabilities of amplitudes.

To evaluate whether voltage-related metrics can explain part of the variance of the calcium transient amplitudes, we performed an analysis at a single event level by selecting all isolated bursts having at least 2 spikes in the burst (n = 650 bursts). The tested metrics were chosen not to be redundant and thus independent of the number of spikes per burst. Thus, we selected the mean frequency corresponding to the average isi within the burst, the duration of the depolarization (the sigma of a Gaussian fit of the depolarization) divided by the burst size, and the mean depolarization amplitude taken at spike onset. Calcium transient amplitudes were normalized per cell and burst size to only probe correlation to the inter-event variability. Linear correlations were performed per cell and Bonferroni correction was applied to account for multiple testing. We then performed partial correlations per cell to see if the identified significant pairwise correlations resist the high interdependencies that exist between voltage-related metrics.
